# Evolution of reading and face circuits during the first three years of reading acquisition

**DOI:** 10.1101/2022.05.14.491924

**Authors:** Xiaoxia Feng, Karla Monzalvo, Stanislas Dehaene, Ghislaine Dehaene-Lambertz

## Abstract

Although words and faces activate neighboring regions in the fusiform gyrus, we lack an understanding of how such category selectivity emerges during development. To investigate the organization of reading and face circuits at the earliest stage of reading acquisition, we measured the fMRI responses to words, faces, houses, and checkerboards in three groups of 60 French children: 6-year-old pre-readers, 6-year-old beginning readers and 9-year-old advanced readers. The results showed that specific responses to written words were absent prior to reading, but emerged in beginning readers, irrespective of age. Likewise, specific responses to faces were barely visible in pre-readers and continued to evolve in the 9-year-olds, yet primarily driven by age rather than by schooling. Crucially, the sectors of ventral visual cortex that become specialized for words and faces harbored their own functional connectivity prior to reading acquisition: the VWFA with left-hemispheric spoken language areas, and the FFA with the contralateral region and the amygdalae. The results support the view that reading acquisition occurs through the recycling of a pre-existing but plastic circuit which, in pre-readers, already connects the VWFA site to other distant language areas. We argue that reading acquisition does not compete with the face system directly, through a pruning of preexisting face responses, but indirectly, by hindering the slow growth of face responses in the left hemisphere, thus increasing a pre-existing right hemispheric bias.

**Highlights:**

- Written words and faces activate neighboring areas of the fusiform gyri, but their developmental trajectory is different.
- The growth of word-induced activation in VWFA is primarily due to schooling.
- The growth of face responses is primarily affected by age rather than by schooling.
- Word and face-related areas exhibit distinct functional connectivity even prior to reading acquisition
- VWFA is initially functionally connected with left-hemisphere spoken language areas, and FFA with amygdala and contralateral FFA.

## 1. Introduction

Learning to read is a cultural invention that deeply modifies the brain (Dehaene, Cohen, Morais, & Kolinsky, 2015). The development of specific responses to the learned script in a left fusiform area, called the visual word form area (VWFA), is the major signature of a literate brain, and numerous studies have reported robust correlations between the level of literacy and activations in this region independently of age, scripts, in normal readers as well as dyslexics (Shaywitz et al., 2002; McCandliss, Cohen, & Dehaene, 2003; Cohen & Dehaene, 2004; Richlan, Kronbichler, & Wimmer, 2011; Dehaene et al., 2015; Martin, Schurz, Kronbichler, & Richlan, 2015; Brem et al., 2020; Feng et al., 2020). The VWFA develops rapidly in children, such that an initial activation to the learned script can already be detected after a few weeks of reading instructions (Brem et al., 2010; Lochy, Van Reybroeck, & Rossion, 2016; Dehaene-Lambertz, Monzalvo, & Dehaene, 2018). In all readers, this activation is systematically located within the occipito-temporal sulcus, just lateral to the fusiform gyrus, surrounded by face- and object-responsive regions (Puce, Allison, Asgari, Gore, & McCarthy, 1996; Weiner et al., 2016). This entire region shows late developmental plasticity and develops at a slower pace than the neighboring collateral sulcus (Gomez et al., 2017). The precise location of the VWFA in each individual is thought to be determined, at least in part, by its pre-existing connectivity, particularly to left-hemispheric temporal language areas (Bouhali, de Schotten, et al., 2014; Saygin et al., 2016; Li, Osher, Hansen, & Saygin, 2020) and parietal areas (Yu et al., 2018; Moulton et al., 2019).

In adults, ventral responses to visual categories are spatially organized in a robustly reproducible way, even in illiterate people (Dehaene, Pegado, Braga, Ventura, Filho, et al., 2010): buildings and landscapes elicit preferential responses medially in the parahippocampal gyrus (PPA), bordered externally by a fusiform region responding to faces (FFA), then by an area in the lateral occipital cortex (LOC) responding more to objects than to scrambled images (Haxby et al., 2001; Hasson, Harel, Levy, & Malach, 2003; Downing, Chan, Peelen, Dodds, & Kanwisher, 2006; Grill-Spector & Weiner, 2014; Weiner et al., 2016). Stimulus selectivity is particularly high and well-documented for faces, a category for which dedicated cortical patches with very high specificity are present in adults (Kanwisher, McDermott, & Chun, 1997; McCarthy, Puce, Belger, & Allison, 1999; Grill-Spector, Knouf, & Kanwisher, 2004; Tsao, Freiwald, Tootell, & Livingstone, 2006; Tsao, Moeller, & Freiwald, 2008). In the context of such a reproducible organization, the culture-dependent emergence of the VWFA offers a unique opportunity to explore how new cultural representations exploit the margin of available plasticity and encroach within a mosaic of pre-existing robust specificities (Dehaene & Cohen, 2007a).

The spatial organization of category-specific responses is known to develop slowly during childhood and even adolescence. In infants, a rough medial-to-lateral partition is already observed during the first year of life, with medial responses to places and lateral responses to faces and tools, yet the activations to faces were initially reported to be indistinguishable from those to tools (Deen et al., 2017), but with improved MRI method, a recent study reported activation selective to faces relative to objects in infancy (Kosakowski et al., 2021). Later in childhood, we and others found that fusiform face responses are clearly present, for instance in kindergartners (Cantlon, Pinel, Dehaene, & Pelphrey, 2011; Dehaene-Lambertz et al., 2018), but continue to slowly grow until adolescence (Golarai et al., 2007; Scherf, Behrmann, Humphreys, & Luna, 2007; Grill-Spector, Golarai, & Gabrieli, 2008; Peelen, Glaser, Vuilleumier, & Eliez, 2009; Natu et al., 2016).

Interestingly, other methods than MRI seem more sensitive in detecting specific responses to faces. Behaviorally, infants are strongly attracted towards faces and face-like stimuli relative to well-matched controls (Johnson, Dziurawiec, Ellis, & Morton, 1991), perhaps even in utero (Reid et al., 2017). Electrophysiologically, face-specific ERP components, such as N290 and P400, have been recorded in the first year of life (Farroni, Csibra, Simion, & Johnson, 2002; de Haan, Johnson, & Halit, 2003; Gliga & Dehaene-Lambertz, 2005, 2007) with a right lateralization compatible with the right-hemispheric bias described in adults (see also Otsuka et al., 2007 using NIRS; de Heering & Rossion, 2015; Adibpour, Dubois, & Dehaene-Lambertz, 2017). Note that these methods are sensitive to quite different properties than MRI that might explain their different sensitivity to faces. ERPs record a sum of activity that might come from different sources within, as well as outside, the fusiform gyrus (e.g. from the amygdala, occipital face areas, posterior superior temporal sulcus). Likewise, in Otsuka et al’s NIRS studies, responses over large cortical patches were summed. This rough spatial resolution contrasts with MRI, where local neuronal activity must be large enough within a given voxel to elicit a vascular response, and also (unless the data is analyzed with subject-specific ROIs) reproducible enough at the same location across participants. Altogether, those converging approaches suggest that face responsivity and selectivity are already present in infancy, with a right-hemispheric lateralization, but perhaps based on a greater spatial dispersion and intermingling of neural responses than in adulthood, such that the fusiform specialization for faces continues to grow slowly throughout childhood and adolescence (Golarai et al., 2007; Golarai, Liberman, Yoon, & Grill-Spector, 2010).

What is the situation for reading acquisition? The “neuronal recycling hypothesis” (Dehaene & Cohen, 2007a) proposes that cultural objects such as written words encroach on cortical areas whose prior biological constraints, inherited from evolution, make them suitable for the acquired function, and which possess sufficient plasticity to adapt to it. Thus, reading acquisition encroaches upon a cortical circuit evolved for visual recognition and naming, particularly for high-resolution foveal objects (Hasson, Levy, Behrmann, Hendler, & Malach, 2002; Szwed, Qiao, Jobert, Dehaene, & Cohen, 2014), and may compete with its prior function for face and object processing. The prolonged maturation of ventral visual areas likely facilitates such cultural recycling during childhood. For instance, playing a video game for hours as a child, learning to read music, or doing math, all lead to increases in ventral visual responses to those specific stimuli in adulthood (Amalric & Dehaene, 2016; Mongelli et al., 2017; Gomez, Barnett, & Grill-Spector, 2019). Reading is one of the beneficiaries of this phenomenon, and indeed several studies have reported an impact of reading acquisition, not only on the growth of word responses in the developing left occipito-temporal cortex, but also, importantly, on the nearby and contralateral activations to faces and objects (Dehaene, Pegado, Braga, Ventura, Filho, et al., 2010; Monzalvo, Fluss, Billard, Dehaene, & Dehaene-Lambertz, 2012; Dundas, Plaut, & Behrmann, 2013; Li et al., 2013; Dundas, Plaut, & Behrmann, 2014; Centanni et al., 2018; Dehaene-Lambertz et al., 2018). In illiterate and literate adults, higher reading ability was associated with a small reduction in responsivity to faces in the left fusiform gyrus, thus increasing the right-hemispheric advantage for faces (Dehaene, Pegado, Braga, Ventura, Filho, et al., 2010) (note that Hervais-Adelman et al (2019) did not replicate this result in a distinct population, we examine this point in the discussion). In children, an increased right-hemispheric bias for faces was found in typical 9-year-old readers relative to 9-year-old dyslexic children (Monzalvo et al., 2012; see also Gabay, Dundas, Plaut, & Behrmann, 2017 for similar results). A similar result was found in a within-subject longitudinal study where fMRI activity was compared in the same children scanned at narrow intervals of a few months prior and after reading acquisition (Dehaene-Lambertz et al., 2018). In those cases, however, the right-hemispheric increase in face responses, while positively correlated with reading scores, was not associated with a diminution of face responses in the left-hemisphere. By contrast, Centanni et al (2018) found that the extent of the VWFA was negatively correlated with the extend of the left face fusiform area (FFA) in 6-year-old children with different reading performances (i.e. able to read from 0 to 80 words). Finally, young struggling readers had lower word selectivity in the VWFA, not because of a lack of response to words but because of larger responses to objects (Kubota, Joo, Huber, & Yeatman, 2019). It is important to stress that those cortical changes may occur without necessarily having any measurable behavioral impact – in fact, recent behavioral studies suggest that performance in word and face processing tasks improves steadily during this period, without interfering with each other (Ventura et al., 2013; Kuhn, Gerlach, Andersen, Poulsen, & Starrfelt, 2021; van Paridon, Ostarek, Arunkumar, & Huettig, 2021).

From these results, three different models have been proposed to explain how the VWFA emerges in the visual areas. First, word responses may develop as an “overlay” over other pre-established responses to objects and faces, perhaps making use of orthogonal vector subspaces within the same cortical territory (Ebitz & Hayden, 2021) (overlapping responses model); this view predicts no competition (Hervais-Adelman et al., 2019). A second possibility is that neurons are initially committed to a given role in visual processing, for instance face recognition, and that some have to change their preferred stimuli during reading acquisition; this view predicts that any increase in reading-related responses would imply a corresponding decrease in face responses (pruning model) (Cantlon et al., 2011; Kubota, Joo, Huber, & Yeatman, 2018). Third, words and other categories of visual stimuli may compete for a pool of initially uncommitted neurons; this view predicts that the growth of reading-related responses in the left hemisphere occurs in initially uncommitted cortical regions, but may block the development of face-related responses in the left hemisphere, thus inducing their preferential growth in the right hemisphere and increasing the right-hemispheric lateralization of face responses (blocking model) (Dehaene-Lambertz et al., 2018). As already noted, these hemispheric changes need not have any impact on behavior, but would be solely seen through brain imaging.

To investigate those possibilities, we recently performed a longitudinal fMRI study in which we were able to repeatedly scan 10 children during their first year of school, thus allowing us to probe the responsivity of VWFA voxels prior to reading acquisition (Dehaene-Lambertz et al., 2018). We observed that words invaded a left-hemispheric site that was initially largely uncommitted, only slightly responsive to tools, and lateral to a region that was already face-selective. In those and nearby voxels, neither face nor object responses decreased during reading acquisition. However, we did find that face responses in the opposite, right hemisphere increased faster and were correlated with reading scores (Dehaene-Lambertz et al., 2018, figure 4). Thus, reading acquisition did not seem to prune away any pre-existing initial responses, but rather to block the further development of face-selective responses in the left hemisphere, and induce it to move to the right hemisphere, in agreement with the blocking model.

Here, our aim was to refine our understanding of the spatial organization of the ventral visual areas at the onset of reading acquisition. Our first goal was to study how the development of a preferential response to words correlates with the development of preferential responses to other visual categories (e.g. faces). Second, we aimed to disentangle the effects of maturation (age) and reading acquisition on the spatial organization of ventral visual areas. A major difficulty confronting prior studies is that age tends to be systematically confounded with reading acquisition, thus making it difficult to separate the cultural and maturational causes of brain development. To partially decorrelate age and reading acquisition, we recruited three groups of typical children who varied in age and reading expertise: 6-year-old pre-readers, 6-year-old beginning readers, and 9-year-old advanced readers. Pre-readers and beginning readers were approximately the same age (∼ 6y) but exhibited very different expertise in reading for two reasons. First, inclusion in French public schools is strictly determined by age on the 1^st^ of January, and thus children differing in just a few months of age may end up with a one-year gap in entry in 1^st^ grade, when reading instruction is mandatory in French schools. Second, some children acquire the rudiments of reading earlier, whether at school or in their families. We thus included in the group of beginning readers some children who were not in 1^st^ grade but exhibited early reading skills, as confirmed by the number of words they could read per minute. Thus, we formed two groups of pre-readers and beginners of roughly the same age but with a difference in reading proficiency. The third group, which served as a contrasting point, were 9-year-old with three years of reading experience (advanced readers).

## 2. Material and methods

### 2.1 Subjects

A total of sixty typically developing children (31 boys and 29 girls) were included in the present study. These children could be divided into two groups according to their chronological age (6- and 9- year-olds children) or three groups according to their reading experience (pre-readers, beginning readers and advanced readers).

Children in the younger group (37 children, 59 to 87 months-old, 24 boys and 13 girls) were all following a normal curriculum and were tested at the end of the academic year, between April and October. The difference in reading performance and age was related to the fact that, in France, all children who are born in a given calendar year (January to December) enter first grade. Thus, the oldest children in kindergarten and the youngest in first grade are separated by only a few months of age, but have different reading instruction. We assessed children’s reading level with “Lecture en une minute” (LUM), a standardized list of words that children were asked to read as fast as possible in one minute. Children tested at the end of 1^st^-grade had an average of 8 months of formal reading instruction and were considered beginning readers (n=13). Thanks to variability in early exposure to print (both at school and in the family), we also included six additional children who were at the end of kindergarten but already able to read more than 15 words/mn. Thus, beginning readers comprised 19 children (10 boys and 9 girls) who were 80 ± 7 months-old and could read 19 to 72 words/mn (41 in average). The pre-readers (73 ± 5 months-old, 9 boys and 9 girls) were all kindergartners who did not receive formal reading instruction and read less than 15 words/mn at the end of kindergarten (6 words/mn in average, ranging from 0 to 15). Although we greatly reduced the age difference between pre-readers and beginning readers while keeping an important difference in reading experience, the beginning readers were still 7 months older, on average, than the pre-readers **(see Fig. 1 and Table 1)**. Below, we use a multiple regression approach to further disentangle the variables.

**Fig. 1.**
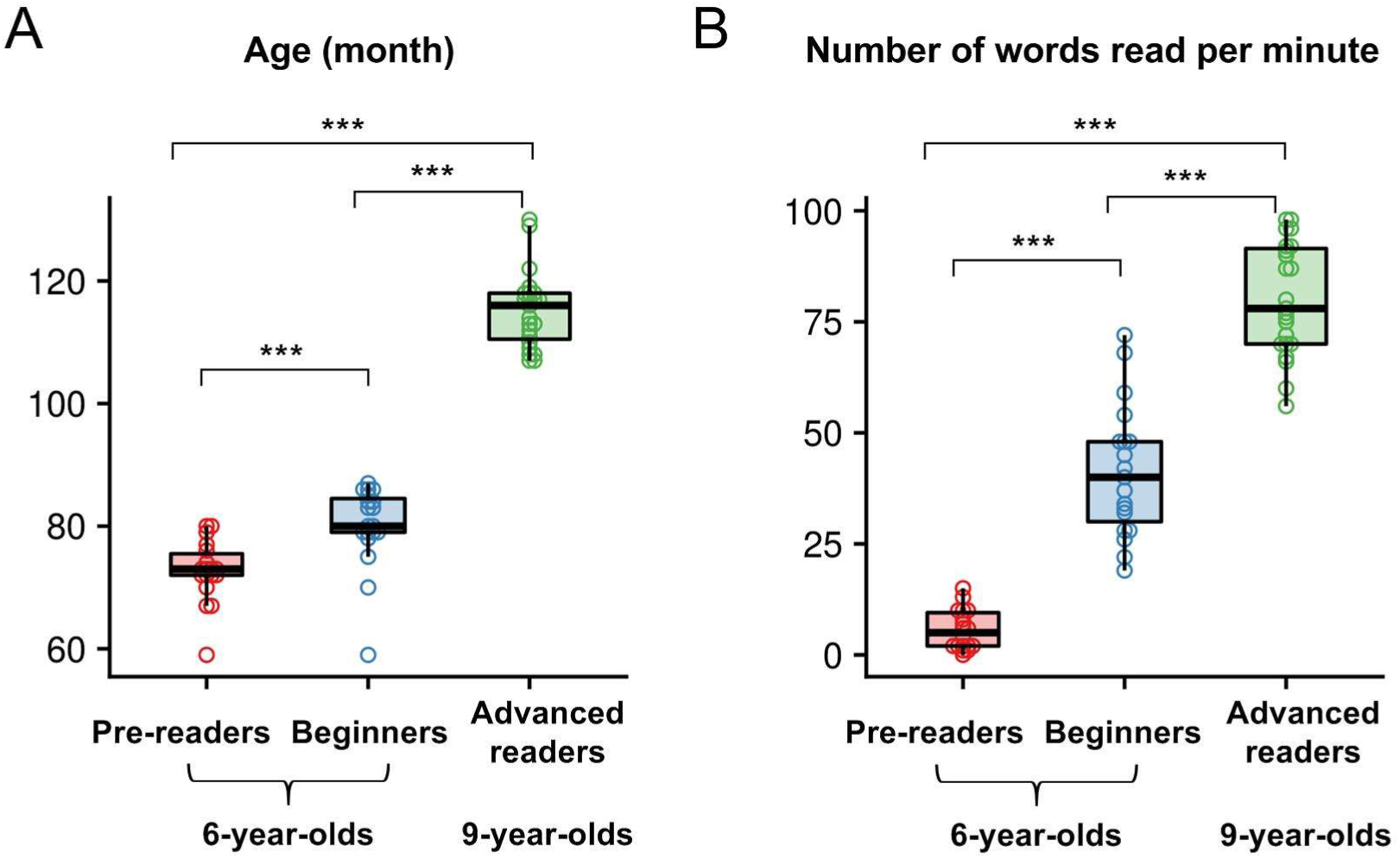
Age distribution (A) and reading performance distribution (B) in pre-readers, beginning readers and advanced readers.

**Table 1.** Characteristics of three groups (mean and standard deviation)

|  | 6-Year-olds |  | 9-Year-old<br>advanced<br>readers | F (2,59) | Pre-readers<br>vs.<br>beginners<br>(p value) | Beginne<br>rs vs.<br>advance<br>d<br>readers<br>(p value) |
| --- | --- | --- | --- | --- | --- | --- |
|  | Pre-readers | Beginning<br>readers |  |  |  |  |
| N | 18 | 19 | 23 |  |  |  |
| Age/months | 73±5 | 80±7 | 115±6 | 295.84** | 0.001*** | <0.001*<br>** |
| Sex | 9M 9F | 10M 9F | 12M 11F |  |  |  |
| Handedness (Left/Right) | 4/14 | 2/17 | 2/21 |  |  |  |
| PIQ<sup>a</sup> (min-max) | 11.62±3.59<br>(4-18) | 10.85±3.78<br>(2-15) |  |  | n.s |  |
| VIQ<sup>b</sup> (min-max) | 10.77±3.27<br>(5-16) | 12.31±1.89<br>(8-15) |  |  | n.s |  |
| Number of words read<br>in 1 min (min-max) | 6±5<br>(0-15) | 41±15<br>(19-72) | 80±13<br>(56-98) | 205.89**<br>* | <0.001**<br>* | <0.001*<br>** |
| Phonological awareness<br>(min-max) | 14.90±9.07<br>(1-28) | 24.95±7.89<br>(0-34) | 30.96±3.95<br>(21-34) | 26.09*** | <0.001**<br>* | 0.025* |
| Rapid automatic naming (min-max) | 25.97±7.32 (16-44) | 20.93±4.82 (14-35) | 16.22±3.16 (12-24) | 17.73*** | 0.014* | 0.015* |
| Number of correctly named images (min-max) | 31.61±4.33 (24-40) | 31.63±8.09 (17-41) | 37.30±5.23 (23-47) | 6.23*** | n.s | 0.011* |
| Verbal short-term memory (min-max) | 13.06±3.31 (8-22) | 13.90±3.86 (8-23) | 18.70±4.74 (12-26) | 11.50*** | n.s | 0.001**<br>* |
| Face memory test (min-max)<sup>c</sup> | 23±2.74 (19-28) | 24±5.41 (15-31) |  |  | n.s |  |

<sup>a,b</sup>: PIQ and VIQ in 6-year-olds are estimated with subtests of the WISC: block design for PIQ estimation, similarities for VIQ estimation. PIQ and VIQ in 9-year-olds were estimated using a different version of WISC test, thus not comparable with the 6-year-olds but all the 9-year-olds have normal PIQ (mean=92.13, SD=16.27) and VIQ (mean=105.22, SD=27.86).
<sup>c</sup>: Face memory test is a subtest of the NEPSY, a neuropsychological battery designed for children aged 3 to 12 years
Note: \* $p < 0.05$ , \*\* $p < 0.01$ , \*\*\* $p < 0.005$

The oldest 9-year-old group (23 children, 115 ± 6 months-old, 12 boys and 11 girls) or the group of advanced reader had three complete years of reading experience (80 words/min in average, ranging from 56 to 98). Their data were previously published in Monzalvo et al (2012).

Children were also tested on several cognitive skills that have been proven to be important for the acquisition of the reading abilities: Phonological awareness was tested with a French standardized test (EVALEC) consisting of the deletion of the first syllable in 10 trisyllabic words, then of the first phoneme in 12 CVC words, and finally in 12 CCV words (Sprenger-Charolles, Colé, Béchennec, & Kipffer-Piquard, 2005). Rapid automatic naming (RAN) of pictures was measured. Vocabulary level was determined with DEN48, consisting in the number (on 48) of pictures correctly named (Jambaqué & Dellatolas, 2000). Verbal short-term memory was assessed with forward and backward digit span and a sentence span (correct repetition of sentences of increasing length).

Pre-readers differed from beginning readers in phonological abilities, rapid automatic naming but not in vocabulary and verbal-short-term memory. Beginning readers differed from advanced readers in all the behavioral tests, including phonological abilities, rapid automatic naming, vocabulary and verbal short-term memory (**Table 1**).

None of the children had a history of neurological, psychiatric or hearing deficit. Their vision was normal or corrected and their IQ estimated by two subtests of the verbal and perceptual domains of the Wechsler’s WISC 3 or 4 was in the normal range. All parents and children gave their written informed consent. The study was approved by the local ethical committee for biomedical research.

### 2.2 Stimuli and experimental task

The stimuli and experimental paradigm were the same as in Monzalvo et al (2012). Four categories of visual stimuli (words, faces, houses, and an expanding checkerboard) were used in a block paradigm. 30 four-letter regular French words and 30 different black and white pictures of unknown people and of houses were used. The words were frequent words encountered by young readers, as specified in Manulex, a lexicon basis comprising occurrence frequency of words in 54 scholar French reading books (Lété, Sprenger-Charolles, & Colé, 2004). Face were front or slightly lateral views of non-famous people with a neutral expression. All stimuli were black on a white background. Faces and houses were highly contrasted black and white photographs matched for size and overall luminance (see Fig 2 for examples). The pictures were the same as in Dehaene et al’s study (2010) in illiterate adults.

**Fig. 2.**
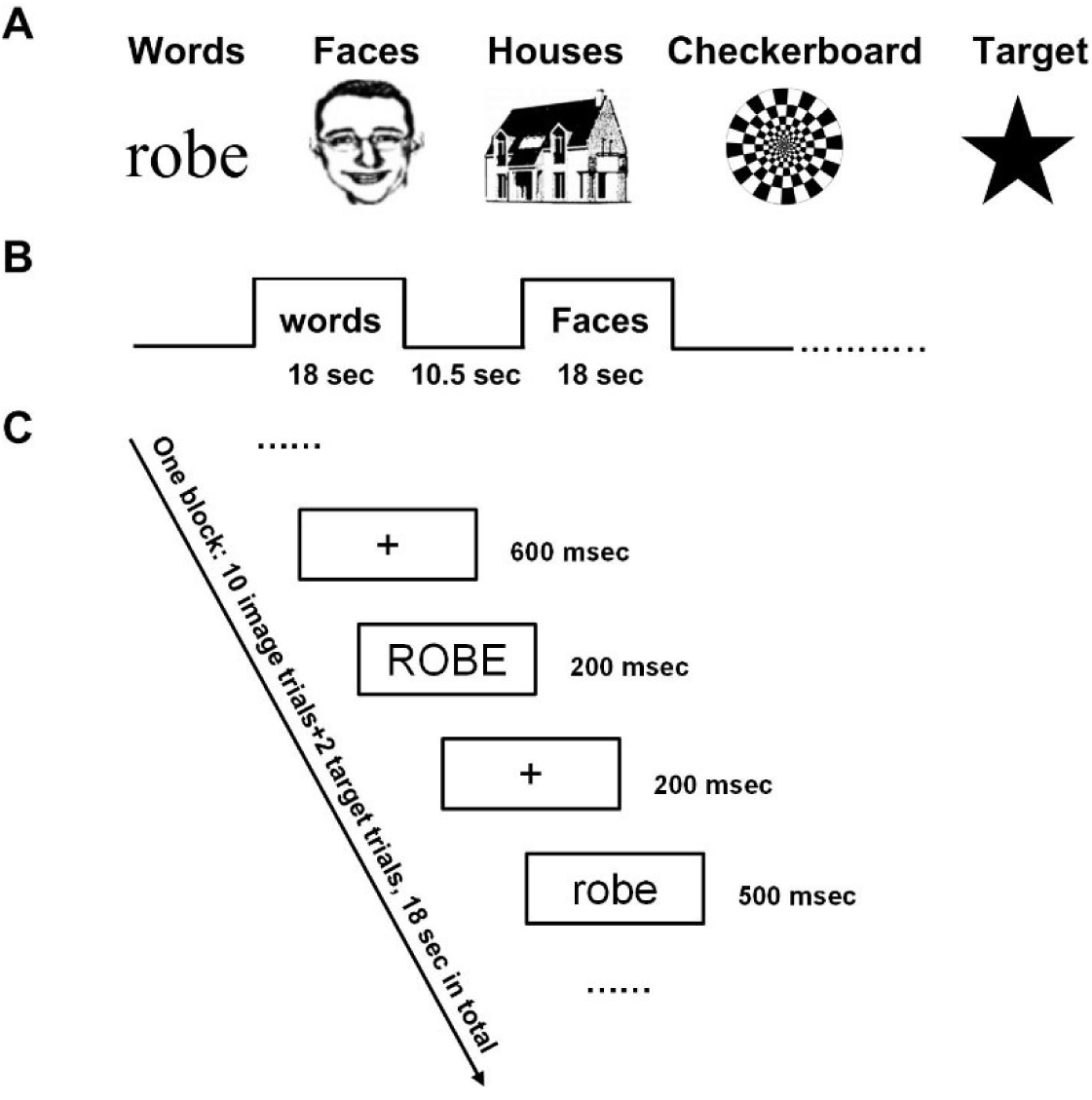
Summary of the experimental design. **(A)** Examples of stimuli used in the fMRI task; **(B)** block design; and **(C)** trial structure of the experiment. In each pair, images varied in size and words had an upper-to-lower case change.

While being scanned, children viewed short blocks of words, faces, and houses and of a looming checkerboard (18 s for each block) followed by a fixation cross during 10.5 s (total bloc duration 28.5 s) (**see Fig.2**). Children performed 4 runs comprising each 7 blocks (two blocks of each category and one block of checkerboards). All blocks were presented in a random order. In each block, 10 pairs of images belonging to the same category were presented (see figure 2; 200 ms presentation for the first item, 200 ms inter-stimulus interval, 500 ms presentation for the second item, 600 ms inter-pair fixation period). On half of the blocks, the images were identical, and on a half of them they differed, but this repetition-suppression design had little effect and is not analyzed here. In addition, two stars (1500 ms duration each) were randomly inserted in each block. Children were instructed to press a button with their left hand whenever a target star appeared. This incidental task was designed to keep the child’s attention toward the visual stimuli.

Before MRI acquisition, each child was trained in a mock MRI scanner, in which the real experiment was simulated using recorded MRI noises and a shortened version of the functional study. They also saw all the words used during the fMRI experiment, in randomly order and case, at their own pace. They were instructed to read what they could. The responses were either the correct word, letters from the word, or “I don’t know”. 500 ms after the child’s response, the following word was presented without feedback. The pre-readers had on average 33.8% correct (whole-word) responses vs 93% and 98.9% for the one-year- and three-year-readers.

### 2.3 fMRI acquisition parameters

Magnetic resonance was performed on a Siemens Tim Trio 3.0 Tesla scanner. Stimulus presentation and behavioral response collection were done with E-prime. Children were protected with noise-protection ear-phones and a mirror system above the child’s head allowed them to see the visual stimuli presented on a screen at the end of the tunnel. T1 images were acquired for anatomical reference (TR = 2300 ms, TE = 4.18, voxel size = 1×1×1 mm, matrix 256×256×176). During the structural acquisition, children were looking at cartoons. For functional imaging, EPI volumes were acquired (TR = 2400 ms, TE = 30ms, matrix = 64×64× 40, voxel size =3×3×3mm). Each visual run comprised 84 volumes.

### 2.4 Data preprocessing

Preprocessing and analyses of the data were conducted using SPM12. The functional images were first corrected for differences in slice-acquisition time and realigned to the first volume in the scanning session. ArtRepair toolbox was used for detecting and repairing bad volumes (43). Two criteria were used to detect bad volumes: (1) 1.5 % variation in the global average BOLD signal from scan to scan, and (2) 0.5 frame-wise displacement, reflecting the sum of all head movements from scan to scan (calculated from realignment parameters). The damaged volumes that exceeded these criteria were replaced by linear interpolation of previous and subsequent images or by nearest-neighbor interpolation when several consecutive images were affected. The framewise displacement (FD), which corresponds to the temporal derivative of the movement parameters was computed for each group and compared. There was a weak but statistically significant difference between the three groups in run 1 (p=0.04), run3 (p=0.017) and run 4 (p=0.047). Post-hoc analyses further revealed that pre-readers moved slightly more than advanced readers (ps=0.036/ns/0.015/0.048 respectively for run 1/2/3/4), whereas there was no significant difference between the two 6-year-old groups in any run, nor between beginners and advanced readers.

For the anatomical image, we first checked for scanner artefacts and gross anatomical abnormalities, then we manually set the origin of T1 image to the anterior commissure for each subject. We normalized each child anatomy to the Montreal Neurological Institute (MNI) template using the DARTEL approach to improve segmentation accuracy and local registration among participants. Functional images were co-registered to their corresponding anatomy. Then the parameters obtained during the DARTEL wrapping process were applied to the functional images which were finally smoothed using a 6 mm Gaussian kernel.

### 2.5 Statistical analyses

#### 2.5.1 Category-specific activations

In each subject, a general linear model was built in which a hemodynamic response function and its time derivative were convolved with block onsets for each category and the 6 motion parameters entered as regressors of non-interest. We implemented a mixed-model analysis of variance (ANOVA) with Group (pre-readers vs. beginning readers vs. advanced readers) as a between-subject factor and Category (Words vs. Faces vs. Houses vs. Checkerboards) as a within-subject factor. We first examined the category-specific activation through the contrasts of one category versus the mean of the other three categories, across the entire set of participants (N = 60), then in each group, and finally compared the 9- and 6- year-olds activations.

#### 2.5.2 Effect of age and reading expertise on brain activity

In order to disentangle effects related to reading from those related to age on brain activity, we conducted two distinct whole-brain regression analyses. First, to investigate the effect of reading acquisition while controlling for age, we focused only on the two groups of 6-year-olds (total n=37) who had approximately the same age but different expertise in reading. We computed analyses with reading performance (number of words read in one minute) as a regressor and with age as a regressor of non-interest. Second, conversely, to examine the effect of age on brain activity, we analyzed the data from all 6- and 9-years-old (total n=60; age range 59-130 months) and entered age as a regressor of interest and reading performance as a regressor of non-interest.

For all analyses, we report effects at a threshold of p < 0.001 at the voxel level and p < 0.05 family wise error (FWE) corrected for multiple comparisons at the cluster level (the cluster-level corrected p value is denoted as p_FWE_corr_).

#### 2.5.3 Number of significant voxels in ventral visual areas

It might be possible that activation is present but more spatially variable or dispersed in younger and/or less expert children, something that would be missed in classical group analyses. To investigate this point, we considered all the voxels showing a preference for one category over the others in a ventral visual mask. The ventral visual mask comprised bilateral inferior temporal gyri and fusiform gyri as defined in the AAL template (Gao et al., 2017; Hervais-Adelman et al., 2019). Category-specific voxels were defined as any voxel in the mask having a z-value > 2.58 (p < .005) for the contrast of this category superior to the mean of the three other visual categories. The number of specific voxels in each child was entered in an ANOVA with Group (pre-readers vs. beginning readers vs. advanced readers) as between-subjects factor, Category (Words vs. Faces) and Hemisphere (Left vs. Right) as within-subject factors. To investigate the effect of reading and age, we also conducted two regression analyses as above: 1) in an analysis restricted to 6-year-olds, the number of words read in one min (LUM) was entered as a continuous predictor variable, while age was the regressor of non-interest; and 2) conversely, in an analysis grouping all 60 children, age was entered as a predictor of interest while reading score (LUM) was a regressor of non-interest.

#### 2.5.4 Medial-lateral and anterior-posterior selectivity

To better investigate the functional organization of the visual cortex, we recovered the activations within 6-mm spheres regularly spaced out along the x axis (x = ±57, ±52, ±47, ±42, ±37, ±32, ±27) at constant y-coordinate (–57) and z-coordinate (–12) on each hemisphere and for each of the categories (words, faces, houses, checkerboard) minus fixation (Dehaene, Pegado, Braga, Ventura, Filho, et al., 2010; Cantlon et al., 2011). The y- and z-coordinates were based on our previous work on illiterates (Dehaene, Pegado, Braga, Ventura, Filho, et al., 2010), children (Feng et al., 2020) and the classic coordinates of the VWFA (Cohen et al., 2002).

The same ROIs approach was also performed along the anterior-posterior axis (y = –80, –68, –54, –42, –30) at the ‘x’ privileged position for words (x= ± 48) and faces (± 39) at constant z coordinate (z = –12) in both hemispheres, since an increasing specificity has been described along this axis (Vinckier et al., 2007; van der Mark et al., 2009). The x and z coordinates were based on the peak of the word- and face-specific activation across all participants.

We first confirmed that we obtained the classic mosaic of preferences for the different visual categories, then we studied the degree of selectivity (selectivity index SI) to words and faces in each ROI and group. We used the simplest definition of SI, namely SI_word_= R_word_ - R(_average of others_) where R_word_ is the BOLD response for words (as estimated by the beta coefficient for words vs. fixation) and R_(average of others)_ is the averaged BOLD response for other categories (Faces vs. fixation, Houses vs. fixation, and Checkerboard vs. fixation). A positive SI denotes a preference for words relative to the other categories, whereas a negative SI denotes a greater response to other categories compared to words.

We computed separate independent-samples t-test among the three groups at each ROI (FDR correction for repeated measures), then conducted our two regression analyses to separate age from reading score, as described above.

#### 2.5.5 Functional connectivity analyses

The present data also offered a possibility to test the biased connectivity hypothesis, according to which the preference for words and faces emergences at specific locations of the ventral occipital-temporal cortex because these locations, even prior to learning, possess privileged connections respectively with the left-hemisphere language network and with the face network (Saygin et al., 2012; Hannagan, Amedi, Cohen, Dehaene-Lambertz, & Dehaene, 2015; Saygin et al., 2016; Li et al., 2020).

We used the spheres defined above as seeds to analyze their “functional connectivity” pattern (i.e. correlation of fMRI signals to other brain sites) and how it varied as a function of reading. The biased connectivity predicted that, even in pre-readers, where selectivity to reading is not yet present, functional connectivity should already be distinct at the VWFA site. To investigate how reading acquisition and age/maturation affect functional connectivity patterns, we focused on anterior-posterior regions near the VWFA and FFA within the occipitotemporal cortex.

Functional connectivity was computed using a seed-voxel correlation mapping analysis (Biswal, Zerrin Yetkin, Haughton, & Hyde, 1995). The ROIs defined above along the posterior-anterior axes and centered on one hand on the VWFA (x = –48, z = –12, y = –80, –68, –54, –42, –30) **(Fig. 6A)**, and on the other hand on the bilateral FFA (x = ±39, z = –12, y = –80, –68, –54, –42, –30) (**Fig. 6D, 6F**) were used as seeds. In each child, we extracted the mean BOLD time series of the voxels comprised in each seed within each word (or face) block, z-scored the signal, and concatenated it across all word (or face) blocks and across the four runs. Three frames (7.2 s) after the end of each task block were included and three frames at the beginning of each task block were excluded in order to account for the hemodynamic delay (Fair et al., 2007). We then computed the temporal correlation between this signal and the similarly processed signal from all the other voxels in a mask defined by the contrast of [(words + faces) > fixation] across all participants. The correlation coefficients were further transformed to Fisher’s Z scores for statistical analyses. Subject-specific contrast images reflecting standardized correlation coefficients were used for the second-level analysis in SPM (Schurz et al., 2015). For each of the five seeds, one-sample t-tests were computed on correlation coefficients to yield functional connectivity maps for each group separately which allowed us to examine seed-specific connectivity (van der Mark et al., 2011).

To further examine the group differences, two-sample t-tests were first computed to determine whether there were reliable group differences in functional connectivity. We then implemented regression analyses with reading performance and age as described above, in order to find regions whose functional connectivity with seeds varied with reading experience independently of age, and vice-versa.

## 3. Results

### 3.1 Behavioral results

Due to unexpected technical problems, the inside-scanner behavioral data of 5 children (2 pre-readers and 3 beginners) were missing. The remaining data showed that 6-year-olds responded less accurately and more slowly to the target star than 9-year-olds. However, there were no significant difference between the two younger groups, neither in the response accuracy (ACC) nor in reaction time (RT) (ACC: main effect of Group: F (2, 52) = 5.48, *p* = 0.007, pre-readers: 79.80% ; beginners: 82.81%; advanced readers: 95.42%; RT: main effect of Group: F (2, 52) = 7.28, *p* = 0.002, pre-readers: 838.35 ± 87.11; beginners: 755.54 ± 113.91; advanced readers: 706.71± 106.30) (see supplementary Table S3). We also verified that the induced activation in the right motor cortex (left-hand response at coordinates [39 –21 60]) was similar in all groups, confirming that all the children were similarly engaged in the task.

### 3.2 fMRI results

#### 3.2.1 Category-specific activations

To determine the local specificity of the activations to the different categories, each category was compared to all others. As shown in **Fig. 2A and 2E**, when all children were analyzed together, the different categories elicited localized activations along a medial-lateral axis in ventral occipito-temporal cortex, from checkerboard in the primary visual and surrounding areas, bordered externally by houses and then faces, and finally words only in the left hemisphere.

Beyond ventral activations, activations to faces were observed in both amygdalae whereas words elicited activations in the posterior superior temporal region, prefrontal regions and angular gyrus in the left hemisphere, and the posterior superior temporal sulcus and angular gyrus in the right hemisphere **(supplementary Table S1)**.

We then examined the responses in each group (**Fig. 2B-D, 2F-H**). A preferential response to checkerboards and houses was seen in each group. Furthermore, as expected, the response to words was present only in the two groups of readers, but not in pre-readers where the closest cluster to VWFA was far from being significant (z-score= 2.14 with p_uncor_=0.016 at [-54, -57, -12]; 4 voxels, p_FWE-cor_=1 and p_uncor_=0.73 at the cluster-level). More surprisingly, the responses to faces also showed striking differences between groups. In the 9-years-old group, the preferential response to faces was firmly established at classical sites in occipital and fusiform regions, together with the bilateral amygdalae **(supplementary Table S2)**. In 6-years-old beginning readers, a specific activation to faces was observed at the same locations. However, in 6-years-old pre-readers, activation in the fusiform regions did not reach significance at the classical p_FWE_corr_ threshold. Only two small clusters were observed at the classical site of bilateral fusiform face areas (left fusiform gyrus: [–39, –51, –21], 5 voxels, z = 3.37, p_FWE_corr_ = 0.96; right fusiform gyrus: [42, –48, –21], 5 voxels, z = 3.61, p_FWE_corr_ = 0.96). Given their low significance, it is impossible to determine whether those findings correspond to false positives or to genuine but feeble activations, whose weakness may also be due to the slightly greater head motion in this group.

**Fig. 2a.**
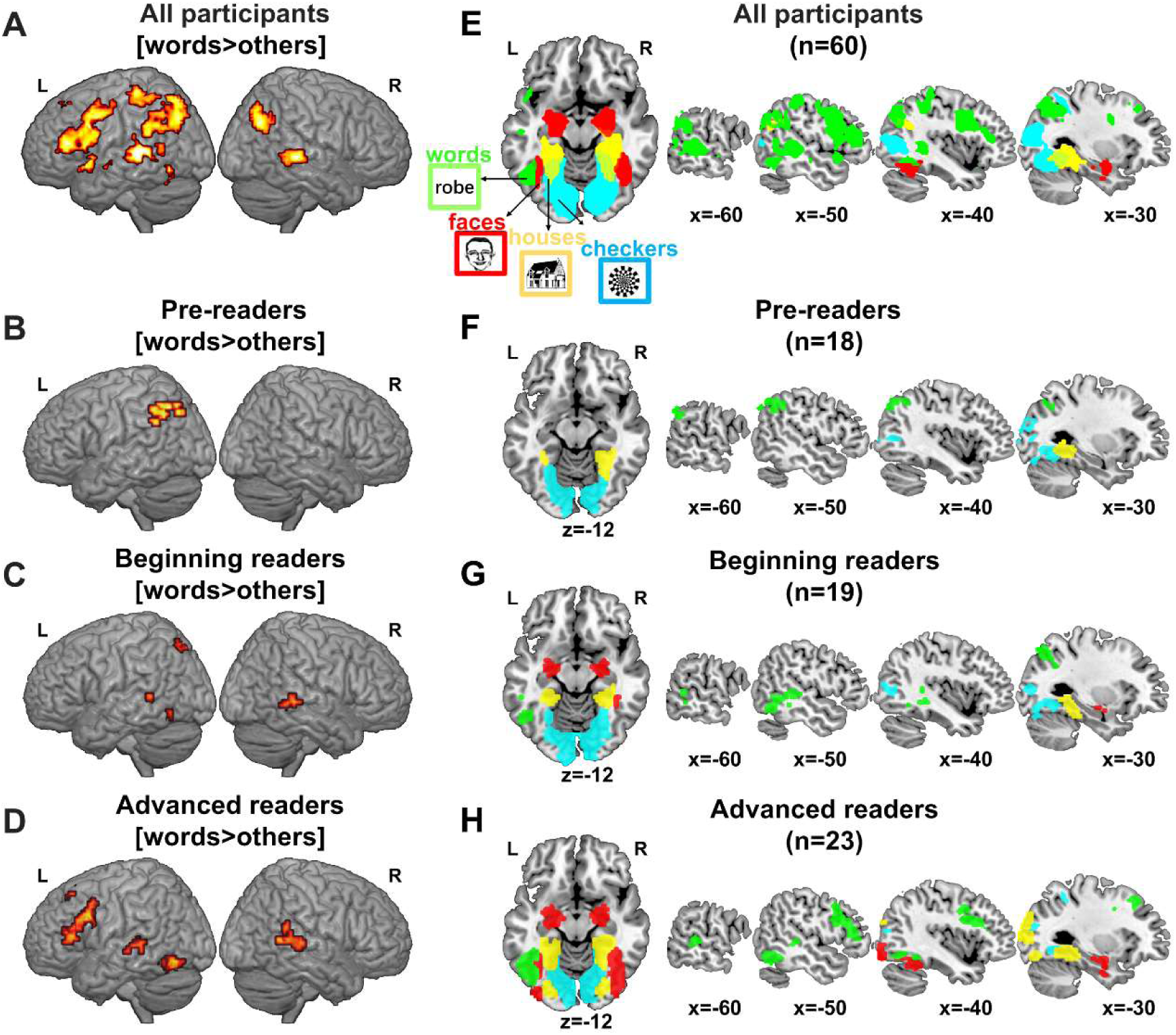
Category-specific circuits. Whole-brain views of words > [other visual categories] for all participants **(A)**, 6-year-old pre-readers **(B),** 6-year-old beginning readers **(C),** and 9-year-old advanced readers **(D)**. Horizontal and sagittal slides showing category-specific activations for all participants **(E)**, 6-year-old pre-readers **(F)**, 6-year-old beginning readers **(G),** and 9-year-old advanced readers **(H)**. Green: regions selectively activated by words (Word > [Face, House, Checker]); Red: face-selective regions (Face > [Word, House, Checker]); Yellow: houses-selective regions (House > [Word, Face, Checker]); Cyan: checkers-selective regions (Checker > [Word, Face, House]). All the regions are reported at the threshold of voxel-level p < 0.001 with cluster-level FWE corrected p < 0.05.

When the 6-and 9-year-old children were compared, no difference in activation to the checkerboard condition was observed, but there was a general increase with age in activation in the right hemisphere for all the other stimuli vs. fixation (words: [27 −75 −6] z = 4.84, 185 voxels, p_FWE_corr_ < 0.001; faces: [27 −69 −5] z = 5.49, 741 voxels, p_FWE_corr_ < 0.001 and houses: [27 −72 −6] z = 5.31, 512 voxels, p_FWE_corr_ < 0.001). There was no significant difference in any of the specific contrasts (one category vs. the others) between the 6- and 9-year-olds.

We then examined the differences between the two 6-year-old groups who had a similar age but very different reading experience. No significant differences were found in the whole-brain analysis. We then focused on the regions with significant activation to a given category in the 9-year-old advanced readers. These regions were used as masks, to which the comparison between the two 6-year-old groups was confined. The only difference was found for words minus others, near the classical VWFA site ([−45 −60 −12] z = 3.39, 8 voxels, p_FWE_corr_=0.05), i.e. the expected result of reading acquisition. There was no significant difference for the other visual categories (Faces, Houses, or Checkerboard vs. other categories)

#### 3.2.2 Regression analyses: Dissociation of the effects of reading expertise and of age/maturation on brain activations

To evaluate the respective effects of reading expertise and age on brain activity to each visual category, we performed two independent whole-brain regression analyses with reading (estimated by the number of words read in 1 min) and age as regressors with one or the other factor cancelled. The goal was to find out which brain regions had activity that varied more with age than with reading level and vice versa.

As expected, specific activations to words (Words > others) correlated with reading level, independently of age, in the left occipitotemporal cortex around the VWFA site when all children were considered (42 voxels, p_FWE_corr_ = 0.02, z = 4.12 at [−45 −63 −9]) but also when only the 6 years-olds were considered (38 voxels, p_FWE_corr_ = 0.03, z = 4.32 at the peak, **Fig. 3A**). No relation between reading level and brain activations to the other visual categories was observed, neither in contrasts such as [one category vs fixation] nor in contrasts [one category vs all others].

**Fig. 3.**
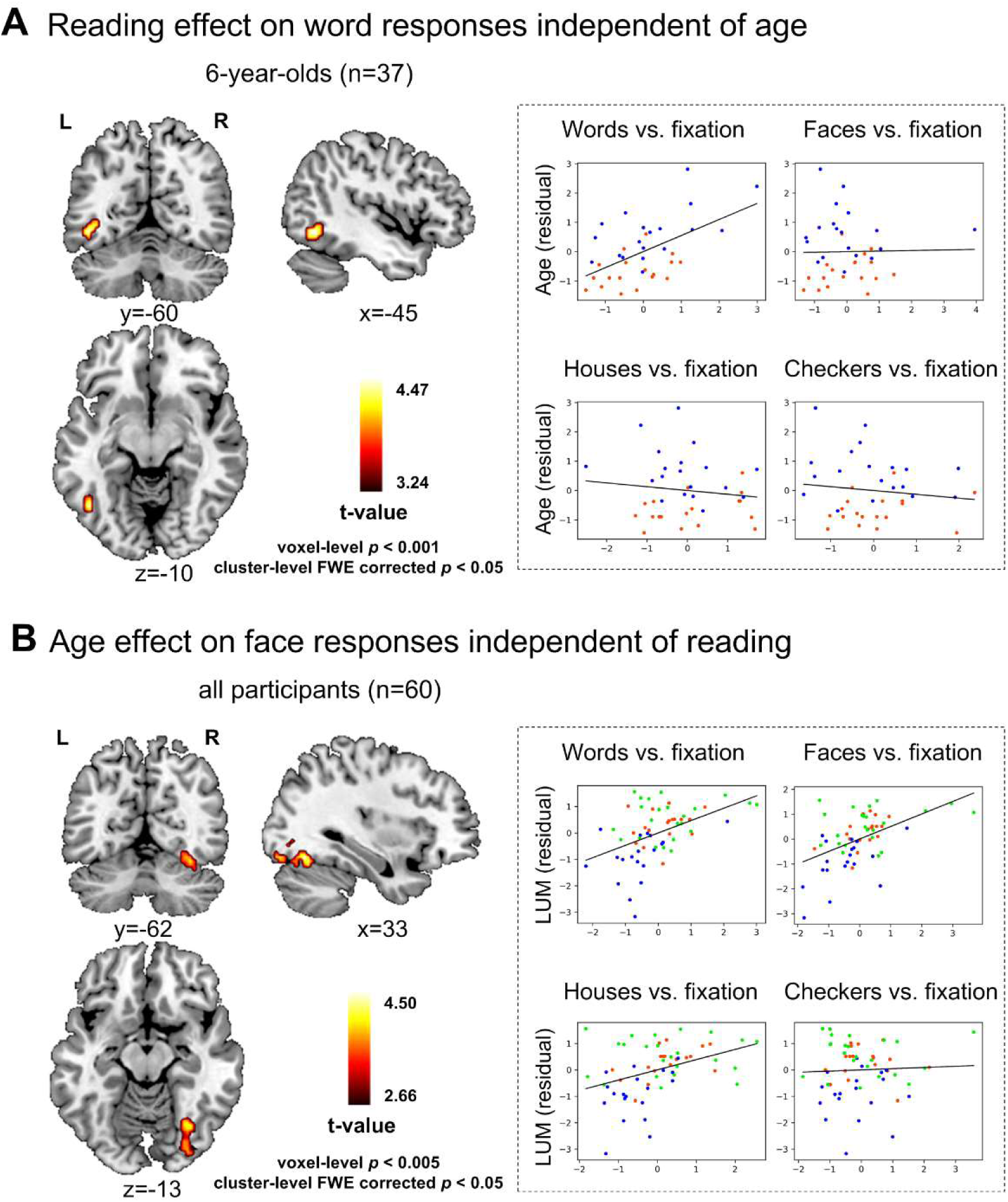
Regression of brain activity with reading and age. (A) left: Effect of reading experience (LUM=number of words read in 1 min) on word-evoked activations relative to others in a model in which age was regressed out in 6-year-olds (n=37). (A) right: Scatterplots of the average activation in the left cluster identified by this analysis for each category (words, faces, houses, checkers) vs. fixation in function of reading, once age was regressed out. (B) left: Age effect on the activation evoked by faces relative to fixation when LUM was regressed out across all children (n=60). (B) right: Scatterplots of the averaged activation in the right cluster identified by this analysis for each category (words, faces, houses, checkers) vs. fixation in function of age, once LUM was regressed out. Each dot, corresponding to a participant is colored in function of the group (pre-readers in red, beginners in blue and advanced-readers in green).

The converse analysis, examining the effect of age independently of reading expertise, revealed a significant cluster in response to faces vs fixation in the right medial fusiform ([33 −66 −15] z = 4.14, 108 voxels, p_FWE_corr_ = .002, **Fig. 3B**) when all children were considered. Note that this cluster was only significant when the voxel-level threshold was lowered to p=0.005 but it survived cluster-level FWE corrected p < 0.05. The same analysis showed no significant cluster when only the 6-year-olds were considered. No significant effect of age on brain activity to other visual categories was observed, neither in the contrast of one category vs fixation, nor the contrast of one category vs all others.

To summarize our results, learning to read allowed the VWFA and the left-hemispheric spoken language network to be activated by written words (**Fig. 2**). While the ventral visual activation to written words was mainly driven by expertise in reading (**Fig. 3A**), the activation to faces was dependent on age (**Fig. 3B**) and was particularly weak in our youngest group. Finally, responses to houses and checkerboards were neither affected by age nor by reading abilities.

#### 3.2.3 Number of category-specific voxels in the visual areas

The above group analyses implicitly assume that there is a reproducible brain localization across children. However, if there was more anatomical variability in the location of the active sites in the younger, less expert children, yet without any difference in activation size, then the group analysis might wrongly conclude to a difference in activation size. To circumvent this difficulty, we also performed a localization-independent analysis on the number of voxels showing a preference for one category (words or faces) over the others in a ventral visual mask. We observed a significant interaction of Group × Category × Hemisphere (F(2, 57) = 4.992, p = 0.01), thus confirming the existence of different selectivity patterns to words and faces in the left and right ventral visual cortex among the three groups (**Fig. 4)**.

**Fig. 4.**
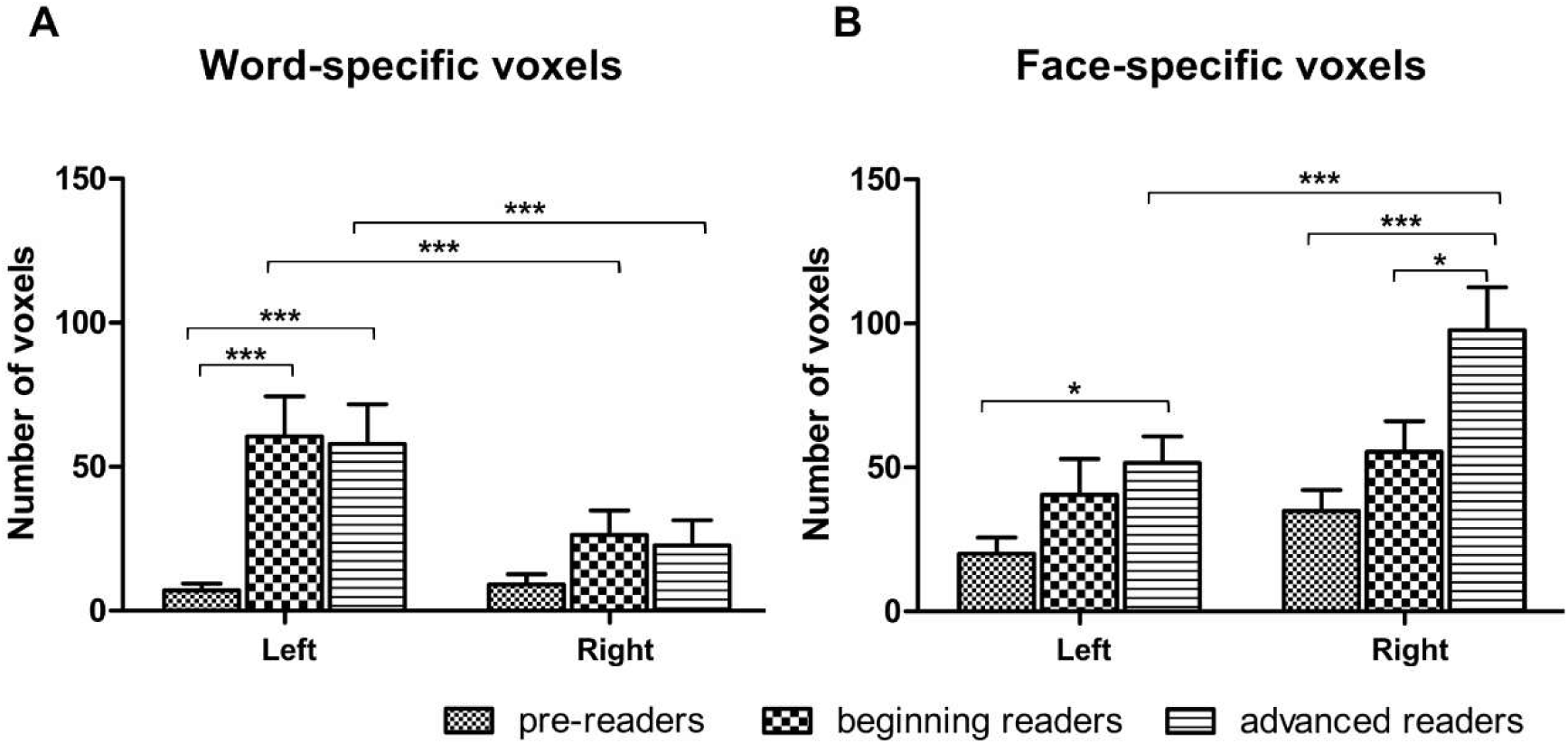
Number of voxels specifically responding to words and faces in the left and right ventral visual cortex in each group. **(A)** Number of word-specific voxels. The number of word-specific voxels in beginning and advanced readers were significantly larger than that in pre-readers and exhibited a significant left lateralization. **(B)** Number of face-specific voxels. The number of face-specific voxels increased with age and became larger in the right hemisphere than in the left hemisphere in advanced readers. *: *p* <0.05; **: *p* <0.01, ***: *p* <0.005

For words, post-hoc analyses revealed that readers (both beginning and advanced readers) had more word-specific voxels than pre-readers in the left ventral visual cortex (beginning vs. pre-readers: *p*_FDR_corr_ = 0.002; advanced vs pre-readers: *p*_FDR_corr_ = 0.003) and also exhibited a left-lateralized pattern with a greater number of word-specific voxels in the left than in the right hemisphere in these two groups (beginning readers: *p*_FDR_corr_ = 0.007, advanced reader: *p*_FDR_corr_ = 0.001). There was no difference in the number of word-specific voxels between the two groups of readers (*p*_FDR_corr_ = 0.89) and no difference between left and right hemispheres in pre-readers (*p*_FDR_corr_ = 0.516) **(Fig. 4A)**. In the 6-year-old children, multiple regression revealed that the number of word-specific voxels in both hemispheres was significantly correlated with reading performance (left: r = 0.522, p = 0.001; right: r = 0.374, p = 0.025) when age was controlled for.

For faces, the number of face-specific voxels in the right hemisphere increased with age: 9-year-old advanced readers had more face-specific voxels than the two 6-year-old groups (advanced vs. pre-readers: *p*_FDR_corr_ = 0.003; advanced vs. beginning readers: *p*_FDR_corr_ = 0.045). However, in spite of a trend for larger face responses in beginning readers compared to pre-readers **(Fig. 4B)**, this difference between the two 6-year-old groups did not reach significance in either hemisphere (left: *p*_FDR_corr_ = 0.121; right: *p*_FDR_corr_ = 0.137). We also observed that 9- year-olds had a right-lateralized pattern with a greater number of face-specific voxels in the right than in the left hemisphere (advanced readers: *p*_FDR_corr_ = 0.004), but this was not the case in the two groups of 6-year-olds (pre-readers and beginning readers, both *P*_FDR_ > 0.1). Interestingly, the number of face-specific voxels in the right hemisphere was also significantly correlated with age when reading performance was controlled for across all participants (r = 0.286 p = 0.028). When restricting the analysis to the 6-year-olds, neither an effect of reading (left: p=0.328, right: p=0.518), nor an effect of age (left: p = 0.839, right: p = 0.449) were found on face-specific voxels in either hemisphere.

No significant difference between groups was found for the number of selective voxels for the house category (**Supplementary Fig. S1**).

#### 3.2.4 ROI analyses of the medial-lateral and anterior-posterior functional organization

Our subsequent analyses were focused on the organization of the ventral visual cortex. Using regularly spaced 6-mm spheres along the x axis (x = ±57 to ±27) at constant y-coordinate (–57) and z-coordinate (–12), we recovered the classic mosaic of preference for the different visual categories along this axis in each group except the pre-readers, who had no word-specific activation at the location of the VWFA (**Supplementary Fig. S2A**). We also performed the same ROIs approach along the anterior-posterior axis [y= –80 to –30] at the ‘x’ privileged position for words (x= 48) and faces (x= 39) in both hemispheres at constant z coordinate (z = –12) (**Supplementary Fig. S2B**). No region anterior or posterior to the classic location of VWFA showed word-specific activation in pre-readers.

In each of the above ROIs, we first compared the selectivity index across the 3 groups (independent-samples t-test), then conducted regression analyses controlling for age and reading performance (**Fig. 5**). The word selectivity index was related to reading performance independently of age around the VWFA [x = (–47), y = –57, z = –12;] **(Fig. 5A)** whereas the face selectivity index was related to age in both hemispheres around the FFA [left: x = –39, y = (– 68), z = –12; right: x = 39, y **=** –54, z = –12] **(Fig. 5D)**.

**Fig. 5.**
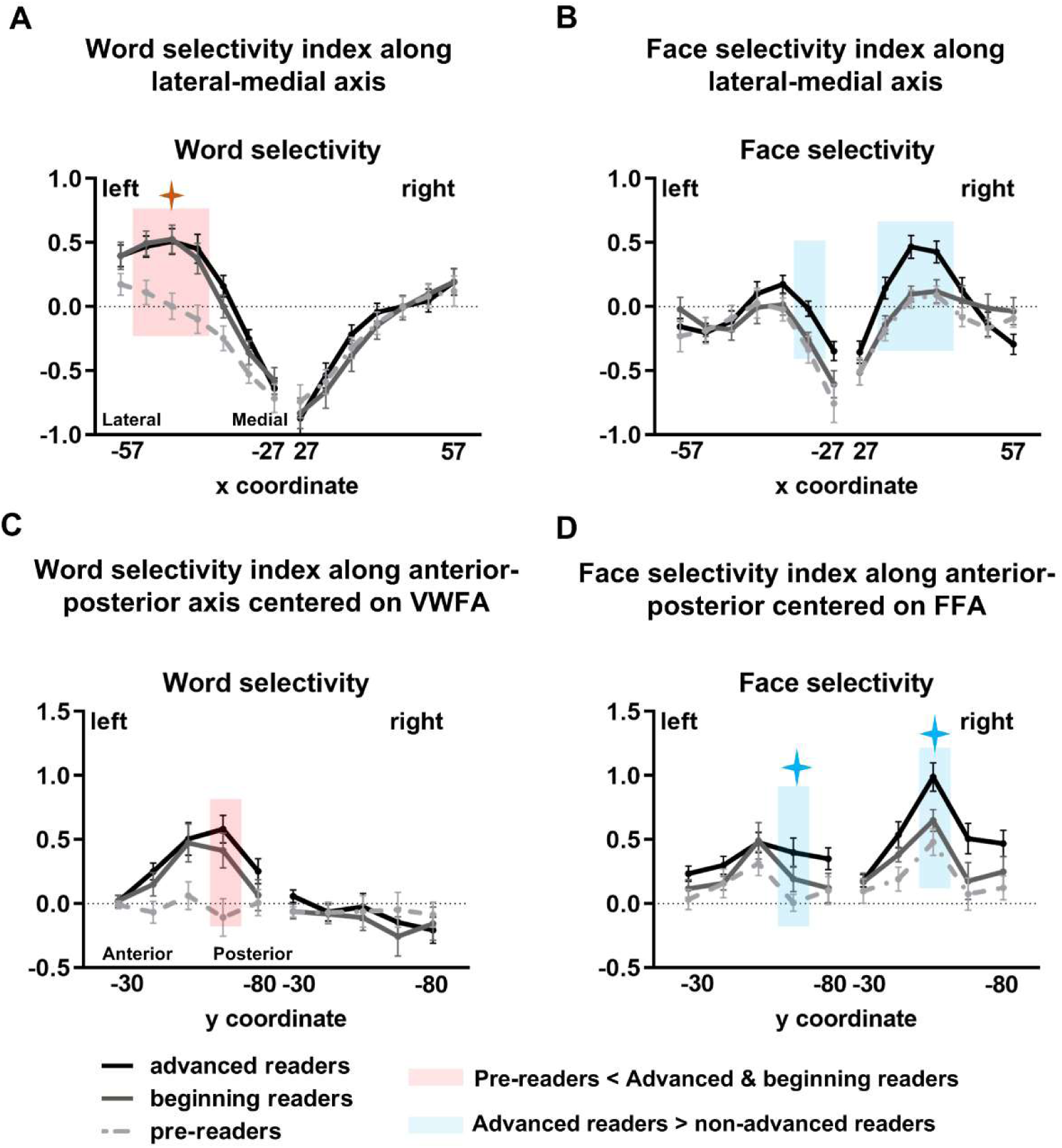
Word- and face-selectivity index in the three groups, along the x axis (A, B) and y axis (C, D). The pink rectangle in the left graphs indicates the locations (in A) [x = (–52, –47, –42); y = –57; z = –12], and in (C) [x = –48; y = (–68); z = –12] where significant differences were observed between pre-readers and each of the two other groups (*p*_FDR_cor_ < 0.05). The blue rectangle in the right graphs indicates the locations (in B) [x = (–32), y = –57, z = –12], and in (D) [x = –39, y = (–68), z = –12] in the left hemisphere; in (B) [x = (32, 37, 42), y **=** –57, z = –12] and in (D) [x = 39, y = (–54), z = –12] in the right hemisphere where significant differences were observed between 9-year-olds and each of the 6-year-old groups (*p*_FDR_corr_<0.05). The red stars further indicate locations in which the word-selectivity index was significantly correlated with reading performance in 6-year-olds when age was controlled; and the blue stars indicate the locations in which the face-selectivity index was significantly correlated with age across all children when reading performance was controlled.

#### 3.2.5 Functional connectivity along the anterior-posterior axis of the fusiform gyri

To test the biased connectivity hypothesis, according to which prior connectivity with other distant cortical regions predicts which sites are likely to become specialized for face and word recognition (Saygin et al., 2012; Hannagan et al., 2015; Saygin et al., 2016; Li et al., 2020), we examined the functional connectivity of the ROIs defined above with all the other voxels activated by words and faces (**Fig. 6**). Along the anterior-posterior axis encompassing the preferred activation to words, all seeds except seed 5 (y = –80) showed significant correlations with left posterior superior temporal gyrus and angular gyrus. Besides, seed 3, which fell closest to the classical coordinates of the VWFA, exhibited the strongest connections with traditional left-hemispheric language areas (left inferior frontal gyrus, inferior parietal lobule and superior temporal gyrus). Crucially, this pattern was significant in all 3 groups, including the pre-readers (see **Fig. 6B**), and we did not observe any significant group differences for any seed in this analysis. In the regression analyses, when we controlled for age and examined the effect of reading performance in 6-year-olds, we found that the functional connectivity between seed 3 and the left middle temporal gyrus increased with reading performance (3 voxels, z = 3.23 at [–60, –36, 9], p<0.001 at peak; see **Fig. 6C**). However, this cluster did not survive correction for multiple comparisons over the whole-brain volume.

**Fig. 6.**
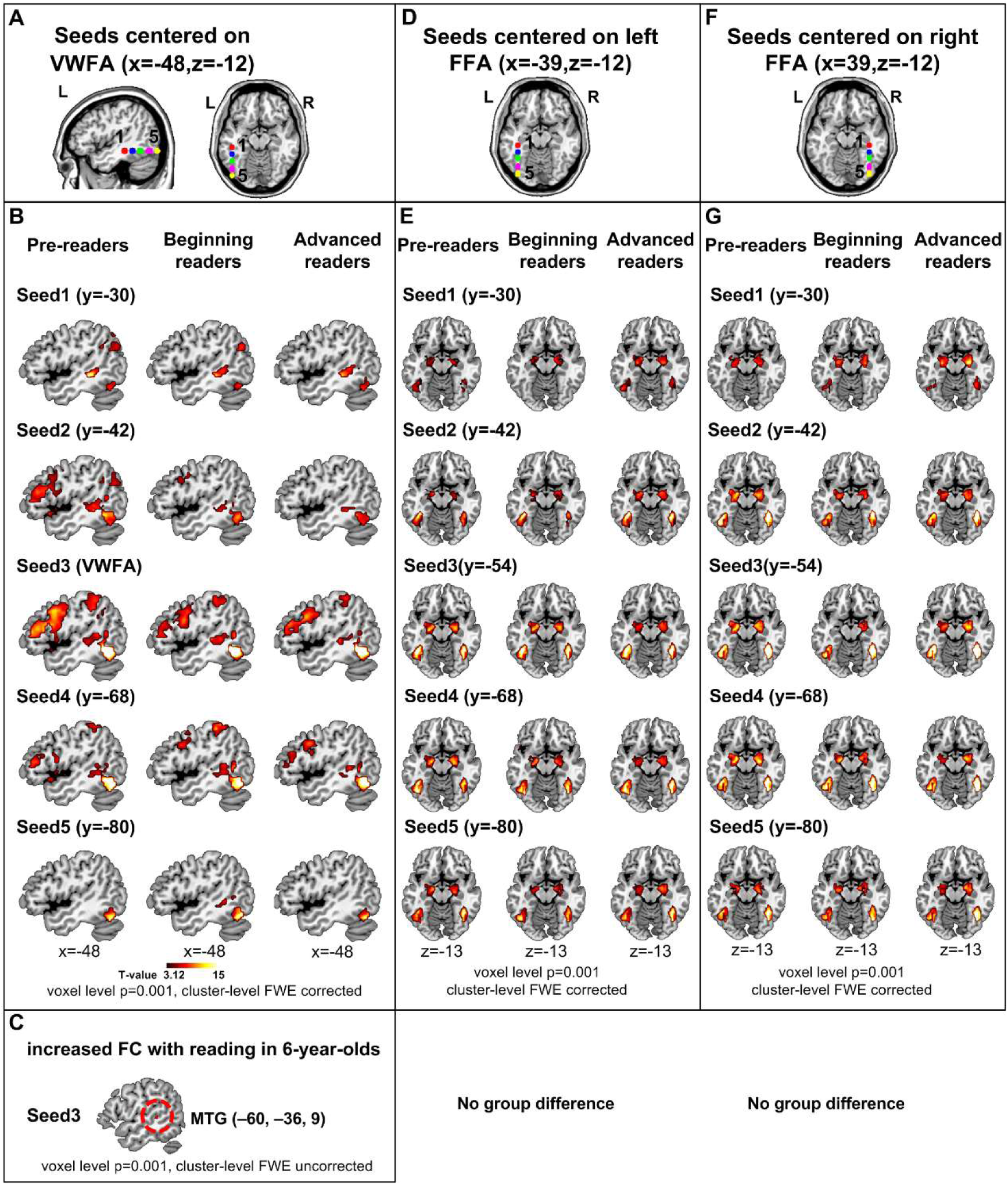
Functional connectivity maps. Illustration of the 5 Seeds in the VWFA-System (x= – 48) **(A)** and FFA-System (x =±39) **(D, F)**: Seed1 (red) was located most anterior, Seed5 (yellow) most posterior in the left occipitotemporal cortex. Seed3 (green) of the VWFA-System corresponds to the center of the VWFA described in previous studies. **(B)** Functional connectivity maps for pre-readers, beginning readers and advanced readers. **(C)** Reading effect on functional connectivity: regions whose functional connectivity with seed 3 positively correlated with reading performance (LUM) in 6-year-olds when age was controlled. **(E, G)** Functional connectivity maps with seeds in bilateral FFA-System for pre-readers, beginning readers and advanced readers.

As concerns faces, the ROIs around the left and right FFAs showed functional connectivity with its contralateral region and bilateral amygdala (left FFA: **Fig. 6E**, right FFA: **Fig. 6G**). We did not observe significant group difference in pair-wise group comparisons. Besides, there was no effect of age on functional connectivity across all children, nor an effect of reading restricted to 6-year-olds.

## 4. Discussion

The human ventral visual cortex displays a robust functional organization, fairly consistent among primates, for natural categories such as faces, bodies, objects and places. Reading is a recent cultural invention based on arbitrary visual symbols that must find its place within this organization. To examine the relative roles of age and reading experience in this transformation, we compared three groups of children of different ages and different reading experiences: 6-year-old pre-readers, 6-year-old beginning readers, and 9-year-old advanced readers. The two 6-year-olds groups differed minimally in age, while their reading levels were distinct and non-overlapping. The difference of a few months of age (∼7 months) that remained between the two groups was further taken into account by including age as a regressor of non-interest in the relevant analyses.

The fMRI results showed that, although written words and faces elicited specific activations in neighboring areas of the fusiform gyri (∼1 cm apart in the left fusiform gyrus, Fig 6), their developmental trajectory was different. On the one hand, a few months of reading instruction were sufficient to observe the emergence of word-specific activations at the classical coordinates of the VWFA. This area was functionally correlated to the spoken language network in all three groups, even before reading acquisition in our pre-readers, and with minimal differences in functional correlation between groups. On the other hand, face responses were affected by age, notably in the right hemisphere. Functional correlations were mainly found with the contra-lateral region and both amygdalae. Finally, the activations evoked by places and checkerboard were affected neither by age nor by reading expertise, and were easily detectable even in the youngest group of pre-readers.

### 4.1 Reading development

Learning to read comprises many stages over the course of several years, each of which could leave its own signature in the brain (Stuart & Coltheart, 1988). According to an accepted developmental hypothesis (Pugh et al., 2001), at the beginning of formal reading instruction, children primarily rely on temporoparietal regions to decode written words in a laborious letter-by-letter manner. Through continuous practice and in tight coupling with the temporoparietal regions, processing in the occipitotemporal VWFA becomes increasingly routinized and unconscious, ultimately resulting in automatic and fluent lexical access and reading performance.

Our findings on pre-readers, beginning readers and advanced readers support this developmental hypothesis. In pre-readers, we found that the contrast of words versus other visual categories only activated the left temporoparietal region. This activation, which lies outside of the ventral visual pathway, fits with the proposal that the processing of written words starts in dorsal attention-related areas (Chyl et al., 2018; Moulton et al., 2019). It might indicate that children without formal reading instruction grope with letters and attempt to guess either their pronunciation (the supra-marginal region being involved in covert articulation; e.g. (Price, 2010)) or to guess words they know by recovering visual and semantic information (Binder, Desai, Graves, & Conant, 2009).

In beginning readers, by contrast, we found that the left temporal-occipital region (VWFA), inferior parietal lobule, left temporal middle gyrus and other regions showed a selective response to words (**Fig. 2**). Even after controlling for age, the VWFA still showed significant differences between the two groups of 6-year-olds, suggesting that the increased activation of this area was caused by children’s reading experience rather than by mere brain maturation. Our results contrast with those of Chyl et al. (2018), who failed to detect VWFA activation in a similar comparison of beginning versus pre-readers (but note their beginning readers could only read 21 words/min on average, compared to 41 for ours). The present finding that VWFA activation starts early on during reading acquisition is, however, convergent with other studies, including our previous longitudinal study which shows a sharp increase of activation in the VWFA after only two months of reading instruction (Dehaene-Lambertz et al., 2018). Furthermore, a previous fMRI training study found that a few weeks of software training sufficed to induce letter selectivity in left occipito-temporal cortex (Brem et al., 2010). The specialization of the VWFA for the learned script is further supported by other studies which found differences in this reading circuit when comparing literate versus illiterates (Dehaene, Pegado, Braga, Ventura, Filho, et al., 2010; Hervais-Adelman et al., 2019) or known versus unknown scripts (Baker et al., 2007; Szwed et al., 2014; Agrawal, Hari, & Arun, 2019). The present research shows that this transformation is mainly due to reading experience, independently of age, as it was only found in those six-year-olds who already showed some reading skills, being able to correctly read on average 41 mots/mn (i. e. approximately one word every 1.5 s).

As expected, advanced readers within 3-years of reading experience activated classical reading areas, including a large activation in left VWFA, posterior superior temporal sulcus and the left inferior frontal gyrus. These activations are similar to adults in a similar task (Dehaene, Pegado, Braga, Ventura, Filho, et al., 2010). They indicate that by 3 years of reading experience, an automatic and efficient access to the language network from visual symbols is already in place.

### 4.2 Reading experience rapidly shapes the word specification in VWFA

While previous studies in children already found that the emergence of VWFA depended on reading experience, this relation was confounded with age. Here, we decorrelated the two effects by comparing two groups of typical children of approximately the same age but with different reading experience. We relied in part on the gap between the continuous variations in children’s age and the sharp definition of which children are allowed to enter school, and in part on the fact that some children acquire rudiments of reading before schooling starts. It should be remembered that even our pre-readers had been exposed to books and some reading/writing in kindergarten, and were able to slowly decipher some of the presented words (∼38% of the words, when tested outside the scanner). Yet our results indicate that their familiarity with print was not sufficient to obtain significant word-specific activation in the VWFA. By contrast, although their reading fluency remained modest (41 words/min in average, half the performance of the advanced readers 80/min), the few months of intense instruction that beginners had received was sufficient to strongly modify this region.

The VWFA is thus one of the most robust and macroscopic evidence of how learning transforms the brain (see also Lochy et al., 2016 using EEG). Brem et al (2010) also reported a fast training effect on brain visual activity in 6-year-old kindergarten children using the *graphogame*, a computer game to teach grapheme-phoneme correspondence. After 8 weeks (3.6 hours), children improved their knowledge but with only slight reading gain. Print-selectivity (Words > False font) increased but in more posterior ROIs than here (i.e. corresponding to our ROIs 4 and 5). These results emphasize the plasticity of the anterior-posterior axis at the lateral level of the VWFA.

The lack of word response in the fusiform region in pre-readers, who have some familiarity with writing may appear in contradiction with Centanni et al (2018), who reported detectable VWFA responses in 44 out of 48 kindergarten children (91.7%). However, in their study, the VWFA was identified in a search area limited to the fusiform area, using a contrast of single letters > faces (rather than letter strings > other categories in our study), and with a lenient criterion (i.e. any voxel with z>2). Furthermore, some of the kindergartners had already developed some reading skills, since their reported word identification skills ranged from 0 to 67 words in the Word ID of the Woodcock Reading Mastery Test (children should read aloud words of increasing difficulty), thus corresponding to a mix of our two groups of pre-reader and beginner 6-year-olds. The use of a one-back task might also have boosted activations contrary to the passive task we used here. Finally, the stimuli differed in size and other visual features, raising the possibility that the activation differences were related to the smaller, more foveal projection of single letters compared to faces, and also to the different spatial frequencies in these two types of stimuli, independently of reading ability (Woodhead, Wise, Sereno, & Leech, 2011). This seems especially likely given the evidence that the VWFA develops in a visual region sensitive to such image properties (Hasson et al., 2002; Brincat & Connor, 2004; Szwed, Cohen, Qiao, & Dehaene, 2009; Gomez, Natu, Jeska, Barnett, & Grill-Spector, 2018). This explanation is plausible given that Centanni et al, identified a right VWFA in 87.5% of children and failed to find any difference between the two hemispheres, in contrast to the systematic left-lateralization of the VWFA, including in beginning readers in the present work (Fig4). Similarly, Dehaene-Lambertz et al (2018) reported a large increase in left VWFA as soon as children entered school and started reading a few words. Using a frequency tagging paradigm in EEG, Lochy et al (2016) also reported a left occipital response to 4-letters non-words rhythmically presented every five items in the middle of false fonts in 5-year-old pre-readers, with an amplitude correlated to the children’s grapheme-phoneme recognition score. Interestingly, familiar symbols (e.g. =,+ !) presented with the same rhythmicity in a control experiment elicited a response on the contralateral right side.

All these results are in agreement with previous proposals that the location of the VWFA is constrained by both visual features such as foveal bias (Brincat & Connor, 2004) or spatial frequency (Woodhead et al., 2011), and by prior connectivity with the language network, thus allowing to associate the arbitrary shapes of letters to phonemes within the left hemisphere. As observed for faces (de Heering & Rossion, 2015), the EEG frequency tagging method might detect the onset of this specialization earlier than MRI, because scattered neurons may add up their activity in an EEG response but may not be sufficiently clustered to induce a BOLD response in fMRI. However, as soon as reading is sufficiently developed for children to read simple words in about one second (our fMRI conditions), regardless of age, our findings indicate that this neuronal specialization becomes large enough to be visible in fMRI.

Finally, we did not observe a larger VWFA activation in 9-year-old advanced readers relative to 6-year-old beginners (fig 4 and 5) although their reading performances were better. This finding might be related to our use of frequent and simple 4-letter words, which were easy to read by both beginning and advanced readers. We might suppose that longer words or more complex graphemes would have elicited larger responses in the better readers. It is also possible that the automatization of reading leads to a progressive reduction and spatial concentration of reading-related fMRI responses, which would go against the initial trend for this response to grow with reading scores and induce a U-shaped pattern over time (for a related idea concerning white matter development and reading, see (Yeatman, Dougherty, Ben-Shachar, & Wandell, 2012).

### 4.5 Functional connectivity of the VWFA and of the FFA

Why do specific and reliable subsectors of ventral visual cortex acquire a specialization for written words? One possibility is that those stimuli differ in ways that fit with pre-existing medial-lateral gradients of preference for features such as line junctions, curvature, foveal retinotopic bias, etc. (Hasson et al., 2002; Szwed et al., 2011; Woodhead et al., 2011; Hannagan et al., 2015). While a foveal bias is now clearly demonstrated (Chyl et al., 2018), visual features alone are clearly insufficient to account for VWFA specialization, because it was found that, when participants were trained to “read” words written in unusual fonts made of faces or houses, those non-letter-like reading stimuli, which normally do not activate the VWFA, can activate it when they are used as reading stimuli (Moore, Durisko, Perfetti, & Fiez, 2014; Martin et al., 2019).

More recently it was suggested that the connectivity to other brain regions might be an important factor in VWFA localization (Hannagan et al., 2015; Saygin et al., 2016). Here indeed, confirming previous anatomical and functional reports (Saygin et al., 2012; Bouhali, Thiebaut de Schotten, et al., 2014; Saygin et al., 2016), we observed that the sites of the FFA and VWFA had very different functional correlations with other brain regions, even prior to reading acquisition. The ROIs most affected by reading experience (x = –48, seed 3) had the strongest functional correlation with the oral language network, even in pre-readers, whereas the more internal seeds at x =± 39, i.e. those with face responses, were all correlated with the contralateral region and both amygdala. It is interesting to observe that in both cases, there was no difference between groups. Notably, the pre-readers who presented almost no selective activation in these ROIs, nevertheless exhibited similar functional correlation with distant areas as the older and more expert children, both in the case of the VWFA and of the FFA seeds. Those findings support the idea that connectivity comes first and drives the localization of functional specialization (Hannagan et al., 2015; Saygin et al., 2016).

Because structural connectivity is largely in place already at birth, a prediction of this view is that similar connectivity biases, predating the face and reading circuits, should already be observed in infants. The scarce data available so far support this view: lateral ventral occipito-temporal regions show a greater functional connectivity at rest with distal associative regions in the temporal and frontal lobes, whereas mesial regions are preferentially connected with ventral and internal areas, including the amygdala (Barttfeld et al., 2018; Li et al., 2020).

All these results suggest that the connectivity to distant regions is an important factor in determining the computational role of ventral visual subregions.

### 4.3 Face-specific activations increase with age

The rapid change in responsivity to written words stands in contrast with the slow development of the face-specific response, whose activation increased between 6 and 9 years **(Fig. 2-5)**, notably in the right hemisphere. Since the first reports of a weak fusiform response to faces in young children (Golarai et al., 2007; Scherf et al., 2007), with a protracted development until adolescence, this observation has questioned models of neural development. Faces are the most frequent and important stimulus with which children are exposed from birth. Even if performance continues to improve until adulthood (Hills & Lewis, 2018), peculiarities of face perception such as the face-inversion effect or the other-race effect already arise during infancy (Sangrigoli & De Schonen, 2004; Hayden, Bhatt, Reed, Corbly, & Joseph, 2007). Infants are even able to link visual facial movements and the produced speech sounds (Weikum et al., 2007; Kushnerenko, Teinonen, Volein, & Csibra, 2008; Bristow et al., 2009; Yeung & Werker, 2013), an ability which In 6-12-y-old children (although not in adults) is correlated with activation in the fusiform gyri and not just the superior temporal sulcus (Nath, Fava, & Beauchamp, 2011).

Given those findings, it could have been thought that face circuitry (including face identification and articulatory movement perception) would have been fully in place prior to reading acquisition. However, many findings, including the present ones, conclusively show that the development of face responsivity in the fusiform gyrus is protracted: fMRI responses selective to faces are detectable in infants (Tzourio-Mazoyer et al., 2002; Deen et al., 2017; Kosakowski et al., 2021) and pre-schoolers (Cantlon et al., 2011), but they continue to develop with age throughout childhood and into adolescence (Golarai et al., 2007; Scherf et al., 2007; Grill-Spector et al., 2008; Peelen et al., 2009).

Here, we reexamined the variables that affect such development. Earlier studies on literate and illiterate adults reported that the growth of the VWFA is negatively correlated with left-hemispheric face responses (Dehaene, Pegado, Braga, Ventura, Filho, et al., 2010). Here, however, in contrast to Centanni et al (2018), we did not observe a negative correlation between the VWFA size and the left FFA size, which would have been suggestive of direct competition or “pruning”. Rather, we found an increase of face responses related to age independently of reading abilities in the right FFA, and an increase of word responses related to reading independently of age in the left VWFA. Similarly, in a longitudinal study of 10 typically developing children during the first year of reading acquisition, we did not observe any direct competition or “pruning” of other visual categories by the growth of reading-related responses (Dehaene-Lambertz et al., 2018). Selecting the voxels that after a year of school were word-specific, it was possible to examine backward their specificity to other visual categories (tools, bodies, faces, and houses) before reading acquisition. These voxels actually had only a weak responsivity to objects, not faces, and this specificity did not change as their response to words increased (Dehaene-Lambertz et al., 2018). The present study, also based on typically developing children, similarly suggests a lack of direct competition between words and faces in the VWFA, but rather the existence of two parallel anterior-posterior axes of word- and face-selectivity, respectively at x = –52 to –42 (for words) and x = –32 (for faces) **(Fig. 5)** (for converging results suggesting the presence of multiple parallel circuits in monkeys, see (Bao, She, McGill, & Tsao, 2020)). Indeed, as noted above, even in pre-readers, the functional connectivity was quite different for those two sites, even though they were only 1 cm apart (fig 6). Congruent with structural connectivity studies (Saygin et al., 2012; Saygin et al., 2016), they favor the hypothesis that reading and face recognition arise from two different core regions that progressively recruit the surrounding cortex (Golarai et al., 2007), rather than from a single region with a gradient of functional specialization as proposed by Behrman and Plaut (2020).

Thus, the present study does not support the “pruning” model, but remains compatible with either the “overlapping responses” model, according to which reading-related emerge on top of pre-existing biases, without interference, or the “blocking” model, according to which reading invades previously uncommitted cortical territory, and blocks the growth of other responses at that site. The present data cannot distinguish between the latter two possibilities, prior studies support the blocking model at least relative to face expansion, because they consistently found differences in amount of rightward lateralization for faces as a function of reading skills, either when comparing literates and illiterates (Dehaene, Pegado, Braga, Ventura, Nunes, et al., 2010; Pegado et al., 2014), dyslexics versus good readers (Monzalvo et al., 2012; Gabay et al., 2017), or within-first-grade children at one year of interval (Dehaene-Lambertz et al., 2018) (but in the same participants, tools and words overlap without interference). Note that the absence of a correlation between reading and face development, in the present work, is not incompatible with this conclusion. The literature suggests that FFA development is very protracted (Golarai et al., 2007; Scherf et al., 2007; Peelen et al., 2009; Pelphrey, Lopez, & Morris, 2009), unlike the few weeks and months that it takes for the VWFA to appear (Brem et al., 2010; Lochy et al., 2016; Dehaene-Lambertz et al., 2018). As elegantly shown by Golarai et al (2007), the FFA progressively expands with age at the periphery of the initial activations. Using the same type of analyses, Dehaene et al (2010) showed that literacy was not affecting the left FFA peak and its close surrounding but the more distant annuli in which face activation was decreasing with reading performances. In the “blocking” model, faces and words are thus competing for the same regions at the periphery of their initial sites. Therefore, a reading-induced bias in FFA development would only emerge when comparing brains with a large difference in reading experience (within or across subjects). Indeed, reading-related differences in FFA lateralization were observed when comparing adults with a lifetime history of literacy versus illiteracy (Dehaene, Pegado, Braga, Ventura, Filho, et al., 2010), 3^rd^ graders normal versus dyslexic readers with a large gap in reading acquisition (Monzalvo et al., 2012) or, within subjects, the same children across more than a year of reading acquisition (Dehaene-Lambertz et al., 2018). The blocking model predicts little or no reading-induced FFA variations when comparing individuals with similar reading experience, such as the present study, where all children were largely funneled through the normalizing effect of the French schooling system, or other studies comparing literate adults from the same culture (Pinel et al., 2014; Davies-Thompson, Johnston, Tashakkor, Pancaroglu, & Barton, 2016).

In possible conflict with this interpretation, Hervais-Adelman et al. (2019) reported no effect of reading skills on activations of other visual categories in any direction. They compared illiterate adults vs. Devanagari (an Indian alphasyllabic orthographic writing system) literates, recruited from the same Indian villages. They also reported correlations between higher reading ability and more left-lateralized response to faces and houses. While we recognize that those data raise a difficulty, we note that reading performance was poor in all groups. It was established by the number of correctly identified letters presented individually during 5s, with up to 10s to answer, and of words presented individually during 10s, with up to 30s to answer. Even under such lenient conditions, the mean reading score measured at two time-points was only 38 (17 to 46) / 42 (30 to 46) letters and 59 (0 to 86) /68 (10 to 85) words in the “literate” group, suggesting that many of the literates were not fluent readers. In fact, the ranges reported in supplementary materials indicate that some “literate” subjects could read… zero words, and only 17 letters. The range of schooling went from 0 to 12 years (∼5 years on average). Thus, Hervais-Adelman et al.’s “literate” group appears closer to the ex-illiterates who never automatized reading than to the fluent readers in Dehaene et al.’s study (2010). Consistent with this interpretation, the average activations of their “literate” group were similar to those of Dehaene’s poor readers. In particular, no larger response to non-words was observed relative to faces within the left fusiform contrary to what was observed in fluent readers (compare figure 4 in Dehaene et al, 2010 with figure 2 in Hervais-Adelman et al, 2019). Such a weakly developed VWFA would not be expected to affect face development.

Finally, we note that some studies did not report competition between words and faces in adult fluent readers but rather a positive effect of reading acquisition on other aspects of visual processing. For instance, Davies-Thompson et al (2016) found a positive relation between the peak MR responses for the two categories within the left hemisphere and between the left VWFA and the right FFA. Positive effects of reading on house activations were also reported in Dehaene et al (2010) and a general increase of the visual responses in Hervais-Adelman et al (2019). As pointed by all the authors, these positive effects can reflect general factors, such as attention, familiarity with books and 2D pictures, as with tests in general, that cannot be disentangled from local effects. Reading acquisition has also a positive effect on early visual cortex, due to the specialization to discriminate small shapes differences, particularly along the horizontal axis, that can benefit other image recognition processes (Dehaene, Pegado, Braga, Ventura, Filho, et al., 2010; Szwed, Ventura, Querido, Cohen, & Dehaene, 2012; Dehaene et al., 2015). Those benefits are not incompatible, and indeed may cooccur with reading-induces changes in the localization and lateralization of visual categories. Future work should endeavor to better separate those two effects.

While a blocking model might explain the expansion of face- and word-specific responses, this hypothesis does not disqualify the possibility of overlapping representations in the same voxels for other categories such as tools, for example. Unlike faces and words, tool and object stimuli are a more heterogeneous category in terms of visual features and function (e.g. shoes vs hammer) which probably accounts for their more distributed response in the lateral occipital cortex and fusiform regions (Downing et al., 2006). In our previous longitudinal study (Dehaene-Lambertz et al, 2018), voxels that become selective to words in the VWFA, also retained their initial selectivity to tools. Similarly, reading expertise generally decreased MRI activations to tools when literate and illiterate adults were compared (Dehaene et al, 2010) but without the distinctive competition pattern observed for faces, which might be related to the distinctive maturation timeline of the fusiform gyrus that we will now consider.

### 4.4 Intense maturation of the fusiform gyri during the schooling period

What makes the ventral visual cortex capable of flexibly acquiring different category specificities throughout childhood? Weiner et al. (2016) investigated the cytoarchitectonic profiles of the ventral visual cortex and described at least four different areas in the fusiform gyri (FG1 to 4) but there was no single association between a cytoarchitectonic area and a unique visual category. For example, area FG4 comprises voxels responsive to bodies, faces and words. In a recent study, Gomez et al (2019) explored gene expression in visual cortices and discovered two series of genes with opposed hierarchical gradients, one ascending and one descending, from posterior to anterior areas comprising ∼200 genes. The authors propose that these gradients signal changes in the distribution of subtypes of neurons synaptic regulation and myelination processes along the visual hierarchy. Age influences the magnitude of gene expression in low- and high-level visual areas along the two gradients, an adult-like pattern being reached around 5 years but with noticeable changes continuing to be present throughout childhood. Furthermore, Gomez et al (2017), by comparing 5-to-12-year-old children to young adults, showed that maturational changes were more intense in the face-selective regions than in the place-selective regions. These structural changes are consistent with what we observed here in a narrower age range (6- to 9-year-olds): responses to faces in the bilateral fusiform gyri increased with age whereas responses to houses were stable. As the task was identical in all blocks (detect the rare occurrence of a target star), the differences between groups and visual categories may indicate genuine difference in the speed and amount of plasticity with different categories are acquired. We could also exclude the possibility that face responses were more dispersed in younger children, because the result remained similar when we considered the number of face-specific voxels in a ventral mask **(Fig. 4)**.

Gomez et al (2017) also reported a decrease in T1 relaxation time between children and adults in the FFA. This finding is thought to reflect microstructural proliferation, notably the growth of the dendritic arborization, that might increase the functional responses by pooling neurons over larger space and/or by refining the specificity of the response through lateral inhibition. All these results emphasize the intense maturational changes that occur in the fusiform gyri in school-age children, precisely when new visual categories such as letters and numbers are intensively learned. They substantiate the neuronal recycling model (Dehaene & Cohen, 2007b), according this architecture is at once prespecified for visual recognition and yet sufficiently flexible to acquire new culturally defined categories. As previously proposed (Dehaene-Lambertz et al., 2018), the newly acquired visual categories may take advantage of this plasticity to invade the ventral visual cortex at a moment where it is still weakly specialized cortex, without having to prune away previously stabilized cortical responses. The situation is very different when adults learn to read: in adulthood, plasticity is much reduced, and the visual categories already in place would strongly compete with the encroachment of a new category for written words (Dehaene, Pegado, Braga, Ventura, Filho, et al., 2010). This may explain why adult learners often struggle to become fluent readers. In the study by Dehaene et al (2010), ex-illiterates were slower to read a list of pseudo-words than were literates. After a 6-month literacy program ∼2 hours per day), the reading performances of the Indian adults remain poor (word reading 0 to 33, mean ∼8 words) and the activation correlated with reading improvement was not at the localization of the VWFA but in occipital areas bilaterally and in the left pars triangularis (Hervais-Adelman et al., 2019). Similarly, adults with pure alexia due to a brain lesion in the left fusiform gyrus have difficulty recruiting the right fusiform as they attempt to reacquire reading (Cohen, Dehaene, McCormick, Durant, & Zanker, 2016) and often remain deficient. Conversely, in a single-case of a child with a large surgical removal of the left ventral occipito-temporal cortex, reading acquisition was easy and a VWFA emerged in the right occipito-temporal sulcus, at a location exact symmetrical location to its usual coordinates (Cohen et al., 2004). The present results confirm that important changes occur in left and right ventral occipito-temporal areas, and demonstrate that some (word recognition) dependent on education while others (face recognition) vary primarily with age, and therefore, presumably, the slow accrual of evidence about faces.

### 4.5 Limitations and future directions

Several limitations of the present study need to be considered. First, keeping attentional effort and performance constant across various age groups is difficult. We choose a task (i.e. detection of a rare star) orthogonal to the stimuli to avoid any difference in performance between younger and older children and between blocks of different visual categories. We assume, but cannot prove, that this task imposes a light and essentially constant load across ages and categories. It is however reassuring that we did not find group differences in the activation evoked by the houses and checkerboards categories, nor by the motor response, suggesting that the children were similarly engaged in the task.

A second limitation is that we acquired only two developmental time points (at 6 and 9 years of age) and simple words and static black and white faces as stimuli. Future studies are needed to test whether the observed increase in activation of the right FFA is continuous with time, and also whether it is stable and corresponds to a genuine partial shift of the FFA from left to right hemisphere. We cannot exclude that the change is transient, due to the particular maturational calendar of the fusiform gyrus during this period. Although the present comparison at the onset of reading is critical, many other phenomena occur later during development that should be investigated in future longitudinal studies.

## 5. Conclusions and implications

The present cross-sectional study converges with the conclusions of Dehaene-Lambertz et al’s (2018) longitudinal study, as well as many recent studies encompassing a larger age range: the developing specialization of word-and face-related areas of the human visual cortex varies with age, education, and the constraints imposed by the connectivity to distant regions. The main contribution of the present research is to show that this growing specialization can occur at two very different time scales. On the one hand, the development of the reading circuit occurs rapidly, driven by a few months of schooling, and regardless of age, at least around six years of age. On the other hand, the development of the face circuit, in spite of six previous years of massive exposure to faces, is far from being finished at that age, and is primarily driven by age. In both cases, however, this development is made possible by the protracted plasticity of the fusiform regions, as elegantly shown through cyto-architectonic and gene expression analyses by Gomez et al (2019). Reading acquisition, and presumably other design features of our school system such as the rapid acquisition of numerals and other mathematical symbols, takes advantage of this remarkable window of plasticity.

## Acknowledgment

This work was supported by INSERM, CEA, Collège de France, and the Bettencourt-Schueller Foundation. We are grateful to the NeuroSpin support teams for their help throughout this study, and to the children and their parents for their interest and participation in this research.

## Supplementary

**Fig. S1.**
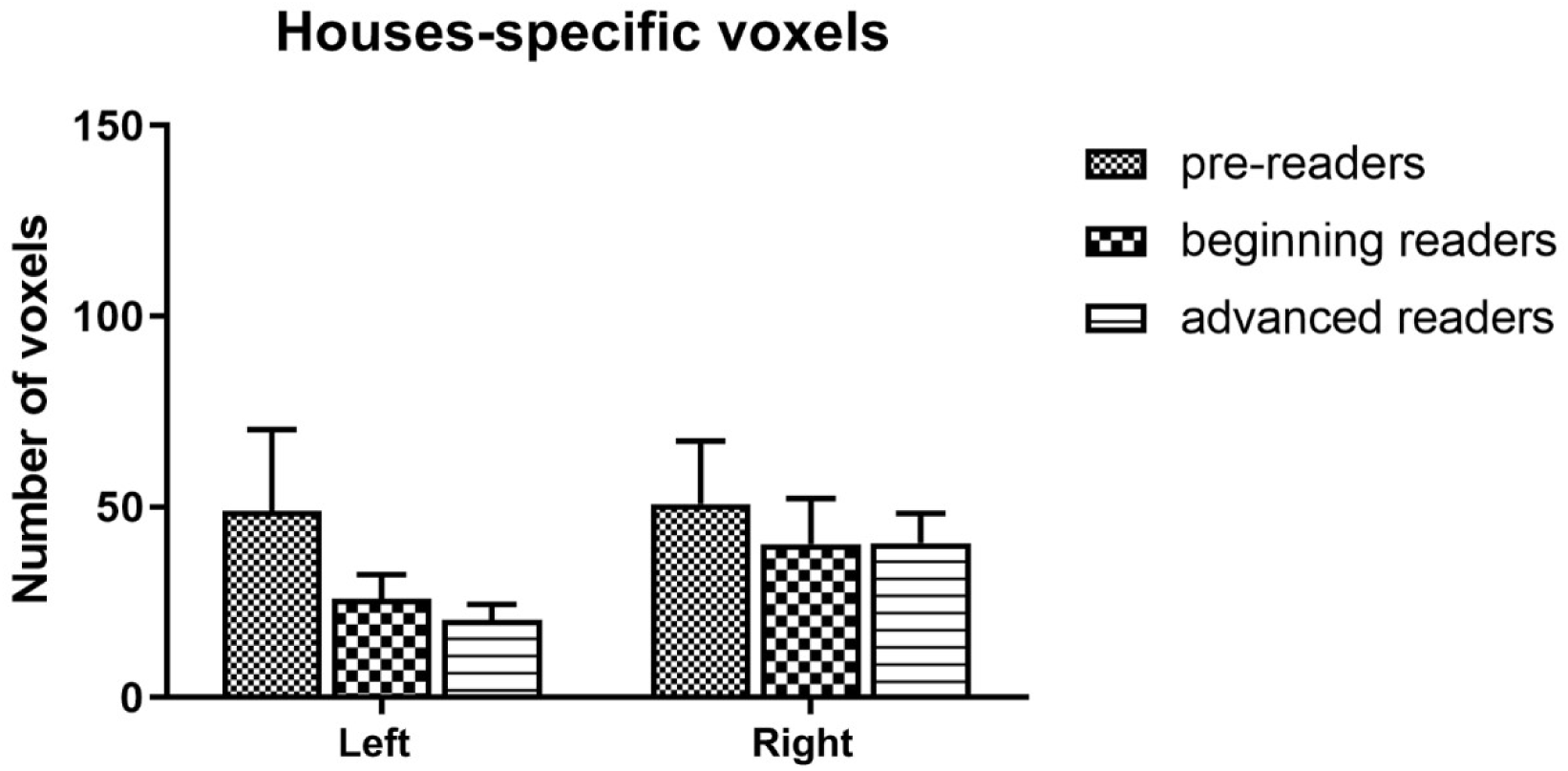
Number of voxels selectively responding to houses in the left and right ventral visual cortex in each group.

**Fig. S2.**
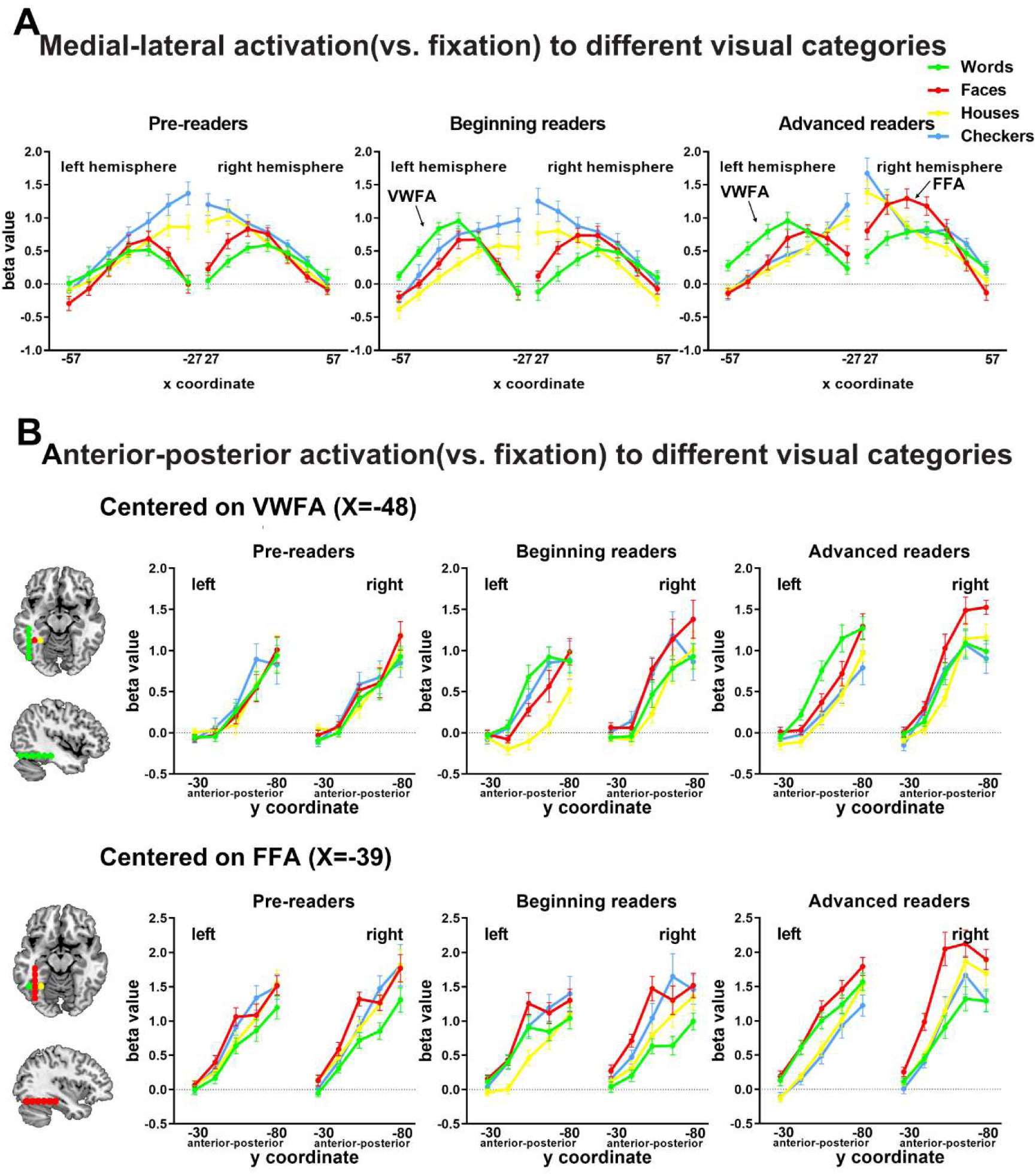
Mosaic of preferences for different visual categories in ventral visual cortex. **(A)** Activation relative to fixation for the different categories in successive spheres of 6 mm regularly spaced at medial–lateral axis (±57 to ±27) in three groups of children (left: pre-readers; middle: beginning-readers, right: advanced readers). **(B)** Activation relative to fixation for different categories in successive cortical sites tracing a vertical line (y-axis) through the classical coordinates of the VWFA (x = –48) and FFA (x = –39).

**Table S1.** Regions of significant activations for each visual category vs. the three others across all participants (individual voxel p=0.001, cluster-level FWE corrected p < 0.05).

| Region | MNI coordinates | | | No. of voxels in cluster | Cluster-level $P$ value (FWE-corrected) | Z value at local maximum |
| --- | --- | --- | --- | --- | --- | --- |
| **Words > others** |  |  |  |  |  |  |
| Left middle temporal sulcus | -60 | -33 | 6 | 474 | <0.001 | 7.40 |
|  | -48 | -39 | 0 |  |  | 6.381 |
|  | -54 | -60 | -12 |  |  | 6.324 |
| Left precentral | -45 | 12 | 30 | 731 | <0.001 | 6.42 |
| Supplementary motor area | -3 | 12 | 54 | 295 | <0.001 | 6.50 |
|  | 0 | -18 | 51 |  |  | 4.16 |
| Posterior cingulate cortex | -3 | -36 | 33 | 278 | <0.001 | 6.34 |
|  | 0 | -33 | 21 |  |  | 5.47 |
| Right middle temporal gyrus | 54 | -36 | 3 | 198 | <0.001 | 5.93 |
| Right angular gyrus | 51 | -69 | 33 | 174 | <0.001 | 5.53 |
| Left angular gyrus | -36 | -66 | 30 | 689 | <0.001 | 5.18 |
| Left Middle frontal gyrus | -27 | 30 | 45 | 109 | <0.001 | 4.45 |
| **Faces > others** |  |  |  |  |  |  |
| Right fusiform gyrus | 42 | -51 | -21 | 121 | <0.001 | >8 |
| Right amygdala/hippocampus | 18 | -9 | -18 | 225 | <0.001 | >8 |
| Left amygdala/hippocampus | -18 | -9 | -18 | 110 | <0.001 | >8 |
| Left fusiform gyrus | -39 | -51 | -18 | 66 | 0.006 | 7.28 |
| **Houses > others** |  |  |  |  |  |  |
| Right fusiform gyrus | 30 | -42 | -9 | 619 | <0.001 | >8 |
| Left fusiform gyrus | -30 | -45 | -6 | 618 | <0.001 | >8 |
| Left middle occipital gyrus | 36 | -87 | 15 | 125 | <0.001 | 5.56 |
| Medial superior frontal gyrus | -6 | 51 | 18 | 250 | <0.001 | 4.72 |
| Left angular gyrus | -42 | -66 | 30 | 86 | 0.001 | 4.25 |
| **Checkerboard > others** |  |  |  |  |  |  |
| Left calcarine | 0 | -84 | 0 | 5442 | <0.001 | >8 |
|  | -6 | -84 | -6 |  |  | >8 |
|  | -15 | -93 | 21 |  |  | >8 |
| Right superior temporal gyrus | 51 | -39 | 15 | 198 | <0.001 | 7.63 |
| Right insular | 33 | 21 | -3 | 141 | <0.001 | 6.55 |
| Right inferior frontal gyrus | 45 | 9 | 30 | 74 | 0.003 | 4.09 |

**Table S2.** Regions of significant activations for each visual category vs. the three others in each group (individual voxel p=0.001, cluster-level FWE corrected p < 0.05).

| Region | MNI coordinates | | | No. of voxels in cluster | Cluster-level $P$ value (FWE-corrected) | Z value at local maximum |
| --- | --- | --- | --- | --- | --- | --- |
| **Pre-readers** |  |  |  |  |  |  |
| **Words > others** |  |  |  |  |  |  |
| Inferior parietal lobule | -48 | -60 | 45 | 108 | <0.001 | 4.15 |
| **Faces > others** |  |  |  |  |  |  |
| **Houses > others** |  |  |  |  |  |  |
| Left lingual | -30 | -45 | -6 | 64 | 0.001 | 4.73 |
| Right middle occipital gyrus | 30 | -90 | 9 | 88 | <0.001 | 4.40 |
| Right fusiform gyrus | 30 | -48 | -3 | 138 | <0.001 | 4.11 |
| **Checkerboards > others** |  |  |  |  |  |  |
| Left calcarine | 3 | -87 | 0 | 2232 | <0.001 | 6.67 |
| Right superior temporal gyrus | 48 | -39 | 15 | 46 | 0.007 | 4.47 |
| Left middle occipital gyrus | -39 | -66 | 6 | 63 | 0.001 | 4.26 |
| **Beginning readers** |  |  |  |  |  |  |
| **Words > others** |  |  |  |  |  |  |
| Right middle temporal gyrus | 57 | -42 | -3 | 60 | 0.001 | 5.13 |
| Left middle temporal gyrus | -42 | -36 | -6 | 106 | <0.001 | 4.34 |
| Left fusiform gyrus | -48 | -57 | -12 |  |  | 3.97 |
| Middle cingulum cortex | 6 | -30 | 33 | 63 | 0.001 | 4.18 |
| Left inferior parietal lobule | -30 | -66 | 45 | 45 | 0.008 | 4.13 |
| Inferior frontal gyrus | -60 | 18 | 15 | 34 | 0.029 | 3.83 |
| **Faces > others** |  |  |  |  |  |  |
| Right amygdala/<br>hippocampus | 21 | -9 | -18 | 66 | <0.001 | 5.35 |
| Left amygdala/ hippocampus | -21 | -9 | -18 | 42 | 0.011 | 5.14 |
| Right fusiform gyrus | 39 | -51 | -21 | 64 | <0.001 | 4.87 |
| **Houses > others** |  |  |  |  |  |  |
| Left fusiform gyrus | -30 | -45 | -3 | 97 | <0.001 | 4.72 |
| Right fusiform gyrus | 30 | -42 | -12 | 80 | <0.001 | 4.61 |
| **Checkerboards > others** |  |  |  |  |  |  |
| Left calcarine | 6 | -90 | -6 | 2393 | <0.001 | 6.52 |
| Right hippocampus | 24 | -30 | 0 | 44 | 0.012 | 5.02 |
| **Advanced readers** |  |  |  |  |  |  |
| **Words > others** |  |  |  |  |  |  |
| Left inferior frontal gyrus | -51 | 12 | 27 | 291 | <0.001 | 5.22 |
| Left fusiform gyrus | -45 | -69 | -12 | 118 | <0.001 | 4.65 |
|  | -54 | -57 | -12 |  |  | 4.22 |
| Right middle temporal gyrus | 54 | -33 | 3 | 98 | <0.001 | 4.22 |
| Left middle frontal gyrus | -30 | 30 | 48 | 59 | 0.002 | 4.07 |
| Left middle temporal gyrus | -63 | -33 | 6 | 62 | 0.001 | 3.92 |
| **Faces > others** |  |  |  |  |  |  |
| Right fusiform gyrus | 39 | -54 | -18 | 201 | <0.001 | 5.56 |
| Left fusiform gyrus | -39 | -51 | -21 | 54 | 0.003 | 5.04 |
| Right amygdala/<br>hippocampus | 21 | -9 | -18 | 112 | <0.001 | 4.75 |
| Left amygdala/ hippocampus | -21 | -6 | -18 | 71 | <0.001 | 4.56 |
| **Houses > others** |  |  |  |  |  |  |
| Right fusiform gyrus | 33 | -42 | -9 | 323 | <0.001 | 6.49 |
| Left fusiform gyrus | -33 | -39 | -9 | 191 | <0.001 | 6.19 |
| Right middle occipital gyrus | 33 | -84 | 9 | 164 | <0.001 | 5.35 |
| Left middle occipital gyrus | -33 | -90 | 21 | 108 | <0.001 | 4.79 |
| Left lingual | -27 | -81 | -12 | 39 | 0.022 | 4.80 |
| Middle cingulum cortex | -9 | -42 | 36 | 70 | 0.001 | 4.30 |
| **Checkerboards > others** |  |  |  |  |  |  |
| Left hippocampus | -21 | -27 | -6 | 35 | 0.029 | 7.17 |
| Right lingual | 9 | -75 | 0 | 3461 | <0.001 | 6.56 |
|  | -9 | -75 | 0 |  |  | 6.42 |
| Right middle temporal gyrus | 45 | -54 | 0 | 146 | <0.001 | 5.41 |
| Left superior temporal gyrus | 48 | -42 | 15 | 100 | <0.001 | 4.51 |
| Left superior parietal lobule | -27 | -51 | 51 | 105 | <0.001 | 4.37 |
| Left middle occipital gyrus | -36 | -63 | 6 | 59 | 0.002 | 4.34 |

**Table S3.**
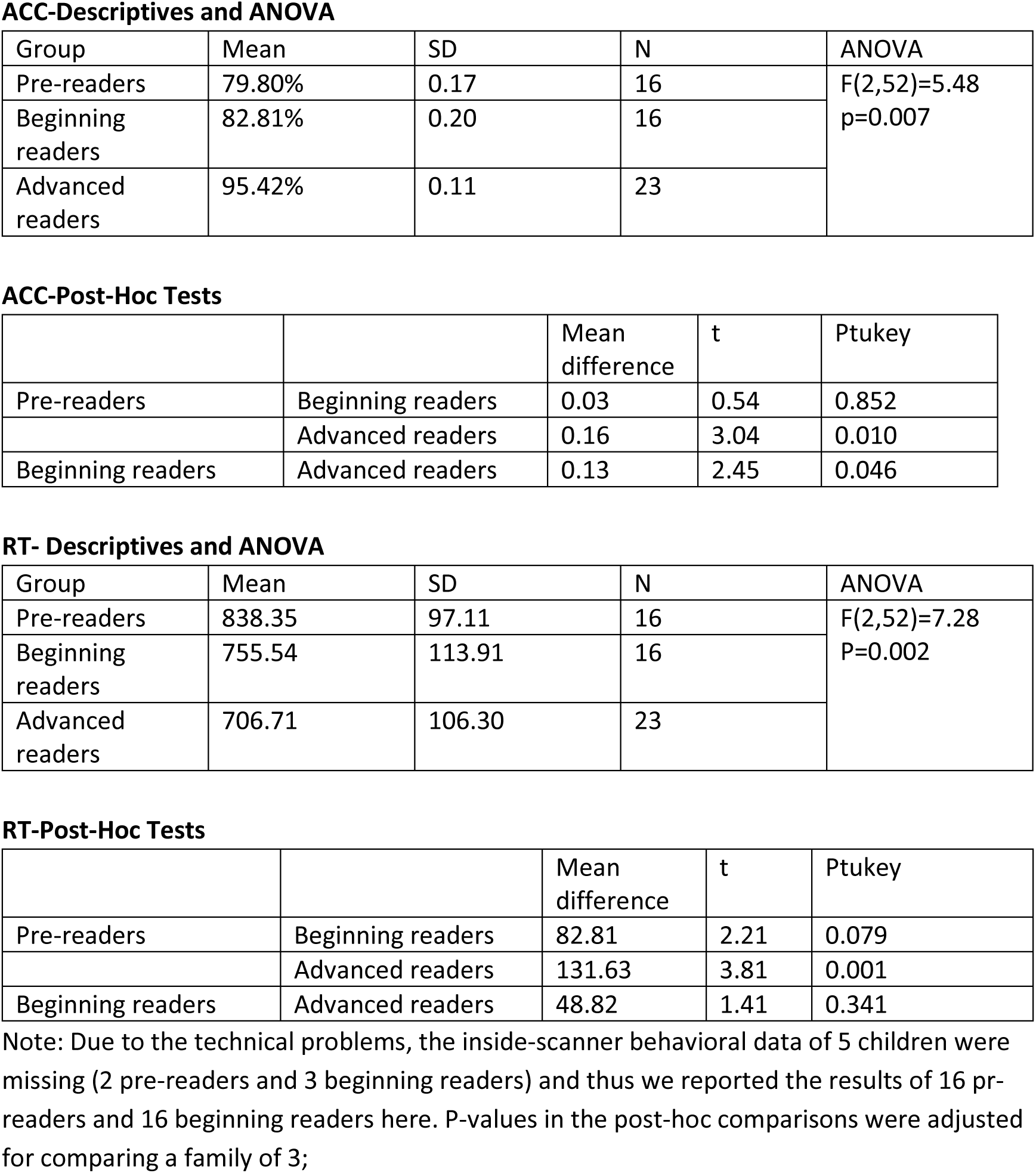
Behavioral performance to the targets in the scanner.

**ACC-Descriptives and ANOVA**
| Group | Mean | SD | N | ANOVA |
| --- | --- | --- | --- | --- |
| Pre-readers | 79.80% | 0.17 | 16 | F(2,52)=5.48<br>p=0.007 |
| Beginning readers | 82.81% | 0.20 | 16 |  |
| Advanced readers | 95.42% | 0.11 | 23 |  |

**ACC-Post-Hoc Tests**
|  |  | Mean difference | t | Ptukey |
| --- | --- | --- | --- | --- |
| Pre-readers | Beginning readers | 0.03 | 0.54 | 0.852 |
|  | Advanced readers | 0.16 | 3.04 | 0.010 |
| Beginning readers | Advanced readers | 0.13 | 2.45 | 0.046 |

**RT- Descriptives and ANOVA**
| Group | Mean | SD | N | ANOVA |
| --- | --- | --- | --- | --- |
| Pre-readers | 838.35 | 97.11 | 16 | F(2,52)=7.28<br>P=0.002 |
| Beginning readers | 755.54 | 113.91 | 16 |  |
| Advanced readers | 706.71 | 106.30 | 23 |  |

**RT-Post-Hoc Tests**
|  |  | Mean difference | t | Ptukey |
| --- | --- | --- | --- | --- |
| Pre-readers | Beginning readers | 82.81 | 2.21 | 0.079 |
|  | Advanced readers | 131.63 | 3.81 | 0.001 |
| Beginning readers | Advanced readers | 48.82 | 1.41 | 0.341 |

Note: Due to the technical problems, the inside-scanner behavioral data of 5 children were missing (2 pre-readers and 3 beginning readers) and thus we reported the results of 16 pre-readers and 16 beginning readers here. P-values in the post-hoc comparisons were adjusted for comparing a family of 3;

